# DePARylation prevents DNA replication-driven PARP1 condensation

**DOI:** 10.64898/2026.08.24.746650

**Authors:** Petar-Bogomil Kanev, Vanya Milanova, Preslava Berkova, Dimitar Kutrovski, Kalina Tosheva, Radoslav Aleksandrov

**Affiliations:** Molecular Mechanisms of DNA Repair Laboratory, Institute of Molecular Biology, Bulgarian Academy of Sciences, Acad. G. Bonchev Str. Bl.21, 1113 Sofia, Bulgaria; BioMedRTC, Institute of Molecular Biology, Bulgarian Academy of Sciences, Acad. G. Bonchev Str. Bl.21, 1113 Sofia, Bulgaria; Institute of Pharmacology and Toxicology, Technische Universität Dresden, Fiedlerstraße 42, 01307 Dresden, Germany

**Keywords:** PARG, PARP1, PARG inhibitor, dePARylation, FUS, biomolecular condensate, FEN1, DNA replication, live-cell imaging, cancer

## Abstract

PARG, the primary enzyme responsible for the reversal of PARP1-mediated poly(ADP-ribosyl)ation, has attracted considerable attention as a therapeutic target in cancer. Yet, the mechanisms underlying PARG inhibitor (PARGi) efficacy remain elusive. Herein, we demonstrate that PARGi prolongs PARP1 residence at damaged chromatin in a manner mechanistically distinct from PARP-inhibitor-induced trapping. That is, PARGi triggers the formation of PAR-driven, FUS-enriched PARP1 nuclear condensates upon DNA damage. Importantly, unrestrained S-phase PARylation during Okazaki fragment maturation also elicited PARP1 condensation in a manner directly reflecting intrinsic PARGi sensitivity, with FEN1 co-inhibition enhancing both PARP1 condensate formation and cytotoxicity. Finally, we discover that dePARylation prevents the rapid nuclear extrusion of PARP1 upon S-phase entry, a phenomenon that is reversible and could undermine PARP1-dependent nuclear processes. Together, our findings reveal that dePARylation precludes the replication-driven condensation of PARP1 and identify Okazaki fragment maturation as a targetable vulnerability that exacerbates condensation and PARGi cytotoxicity.

## Introduction

Chromatin poly(ADP-ribosylation) (PARylation) is a rapid response to DNA breaks, driven primarily by the poly(ADP-ribose) polymerase 1 (PARP1) enzyme and, to a lesser extent, PARP2.^1–3^ PARP1 binds single- and double-stranded breaks (SSBs and DSBs), which triggers structural changes leading to an active conformation, whereby the enzyme can utilize NAD^+^ to produce ADP-ribose units which are attached onto substrate proteins in polymer chains of varying length and branching pattern, termed poly(ADP-ribose) (PAR).^4–6^ The major targets of DNA damage-induced PARylation are PARP1 itself and histones in the vicinity of the breaks.^7^ PARylation decondenses damaged chromatin and serves as a binding platform for a plethora of PAR-binding proteins implicated in chromatin remodeling and DNA repair.^8–11^ Damage-induced PARylation is swiftly reversed by several enzymes, including poly(ADP-ribose) glycohydrolase (PARG), which cleaves glycosidic bonds within PAR, and ADP-ribosylhydrolase 3 (ARH3), which also recognizes glycosidic bonds, but primarily cleaves the proximal Ser-linked ADP-ribose moiety.^12–16^ Importantly, PARG deficiency causes embryonic lethality in mice and severe neurodegeneration in fruit flies, while certain ARH3 mutations cause neurodegeneration in humans, highlighting the importance of efficient PARylation reversal.^17–19^

The pronounced efficacy of PARP1/2 inhibitors (PARPi) in treating homologous recombination (HR)-deficient tumors and the clinical introduction of PARPi compounds has fostered major efforts toward a comprehensive understanding of PARP1 function in genomic stability.^20–22^ Furthermore, the differential cytotoxicity of PARP inhibitors and the capacity of some of these to enhance the binding of PARP1 to DNA breaks (often termed PARP trapping) has put a particular emphasis on PARP1 dynamics at DNA damage sites.^23–25^ Despite conflicting in vitro and in vivo evidence on the matter, our recent live-cell studies provide a coherent model for the mechanism underlying the PARPi-induced PARP1 retention at sites of DNA damage.^23^ Our data revealed that catalytic inhibition of PARP1 results in repeated, unproductive binding-dissociation cycles of inactive PARP1 molecules to DNA lesions. In addition, most PARP inhibitors induce reverse allosteric effects of greatly varying magnitude that prolong the DNA lesion-bound state of PARP1. Both effects combine to determine the overall retention of PARP1 at damaged chromatin sites. The extent of these dynamic shifts strongly correlates with downstream repair delays and PARPi cytotoxicity, highlighting PARP1 retention at breaks as a major challenge to genome homeostasis.^23^

In light of the success of PARP inhibition, targeting dePARylation has also attracted considerable attention as a therapeutic strategy in cancer.^26–28^ As a result, multiple PARG inhibitors (PARGi) have been developed over the past decade, some of which have already entered clinical trials, with a particular focus on breast, ovarian, prostate, and colorectal tumors.^29–35^ А number of genetic vulnerabilities to PARG inhibition have been identified, with studies highlighting overlapping, yet distinct sensitivity profiles when compared to PARPi.^36–39^ PARG inhibition or deficiency has been shown to sensitize cells to genotoxins, including ionizing radiation, topoisomerase I (TOP1) poisons, cisplatin, and alkylating agents.^40–43^ Meanwhile, how DNA repair unfolds within the hyperPARylated environment has remained elusive. Limited evidence suggests that PARG depletion as well as inhibition prevent excessive PARP1 accumulation at damage sites.^44,45^ In contrast, increased PARP1 chromatin retention was reported in cells lacking PARG.^37^ Further corroborating the clinical relevance of PARG, its downregulation has been described as a mechanism of PARP inhibitor resistance.^44^ Rather conflicting, these findings warrant a careful assessment of PARP1 behavior under conditions of unrestricted PARylation.

Herein, we report that PARG inhibition triggers the enhanced accumulation and prolonged presence of PARP1 at complex DNA lesions. Despite an obvious resemblance to PARPi-induced changes in PARP1 dynamics, which included a drastic reduction in turnover at damage sites, this effect was PARylation-dependent and mechanistically distinct, with no consequences for downstream DNA repair events. Extending this dynamic phenotype, which we term “PARP1 capture”, we demonstrate that PARG-driven dePARylation prevents the formation of aberrant PARP1 condensates as a general response to exogenous DNA damage. These condensates exhibited a tendency to originate from PARP1-rich nucleoli and recapitulated the general features of PAR-driven condensate assemblies, including the presence of FET family proteins. Notably, we demonstrate that endogenous PARylation, particularly during S phase, is sufficient to elicit PARP1 condensation in a manner that reflects intrinsic PARGi sensitivity. Amplifying S-phase PARylation through the inhibition of canonical Okazaki fragment maturation greatly enhanced PARGi-driven PARP1 condensation and even forced it in PARGi-resistant cells, sensitizing them and resulting in profound cell death. Lastly, we show that dePARylation prevents the nuclear export of PARP1 during S phase, as PARG inhibition forced a rapid nuclear-to-cytoplasmic translocation of PARP1 upon S-phase entry. This phenomenon was reversible, independent of programmed cell death, and enhanced via FEN1 co-inhibition, with substantial PARP1 nuclear re-entry observed during G1 and G2. Hence, our data establish PARP1 condensation as a biomarker of PARGi sensitivity and Okazaki fragment maturation as a potent intrinsic driver of this phenomenon, while also identifying a previously unrecognized role for PARG and dePARylation in regulating the subcellular localization of PARP1.

## Results

### PARG inhibition prolongs PARP1 residence at DNA damage sites in a PAR-dependent manner

We set out to explore how PARG inhibition influences repair dynamics at sites of DNA damage, to which end we employed our previously described workflow that combines UV laser micro-irradiation (IR) with time-lapse imaging of live cells expressing fluorescently labelled proteins of interest at near-physiological levels.^23,46–48^ Successful PARG inhibition via PDD00017273 (hereafter referred to as PDD^31^ was confirmed based on multifold delays in the removal of PAR degraders PARG (t_1/2_removal=105.27±12.39 s in untreated cells vs. 1973.06±377.56 s in 10 µM PDD-treated cells) (Figures 1A-1C) and ARH3 (103.49±15.82 s vs. 295.81±37.72 s, respectively) (Figures 1D-1F) from damaged chromatin sites, indicative of persistent damage-induced chromatin PARylation. This effect was concentration-dependent, as evidenced by the increasing half-times of ARH3 removal with rising PDD concentrations (Figures 1G and 1H). XRCC1, an archetypal PAR binder and base excision repair (BER) scaffold protein,^9^ also exhibited delayed removal (Figures S1A-S1C), in addition to an enhanced accumulation at lesions (Figures S1A, S1D, and S1E). COH34^29^ and JA2131^30^, two additional PARGi, did not alter PARG (Figures 1A-1C), ARH3 (Figures 1D-1F), or XRCC1 dynamics (Figures S1A-S1E). The prolonged accumulation of PAR-binding factors under PDD treatment confirmed a hyperPARylated chromatin state at damage sites, which the other two PARGi compounds failed to elicit.

**Figure 1.**
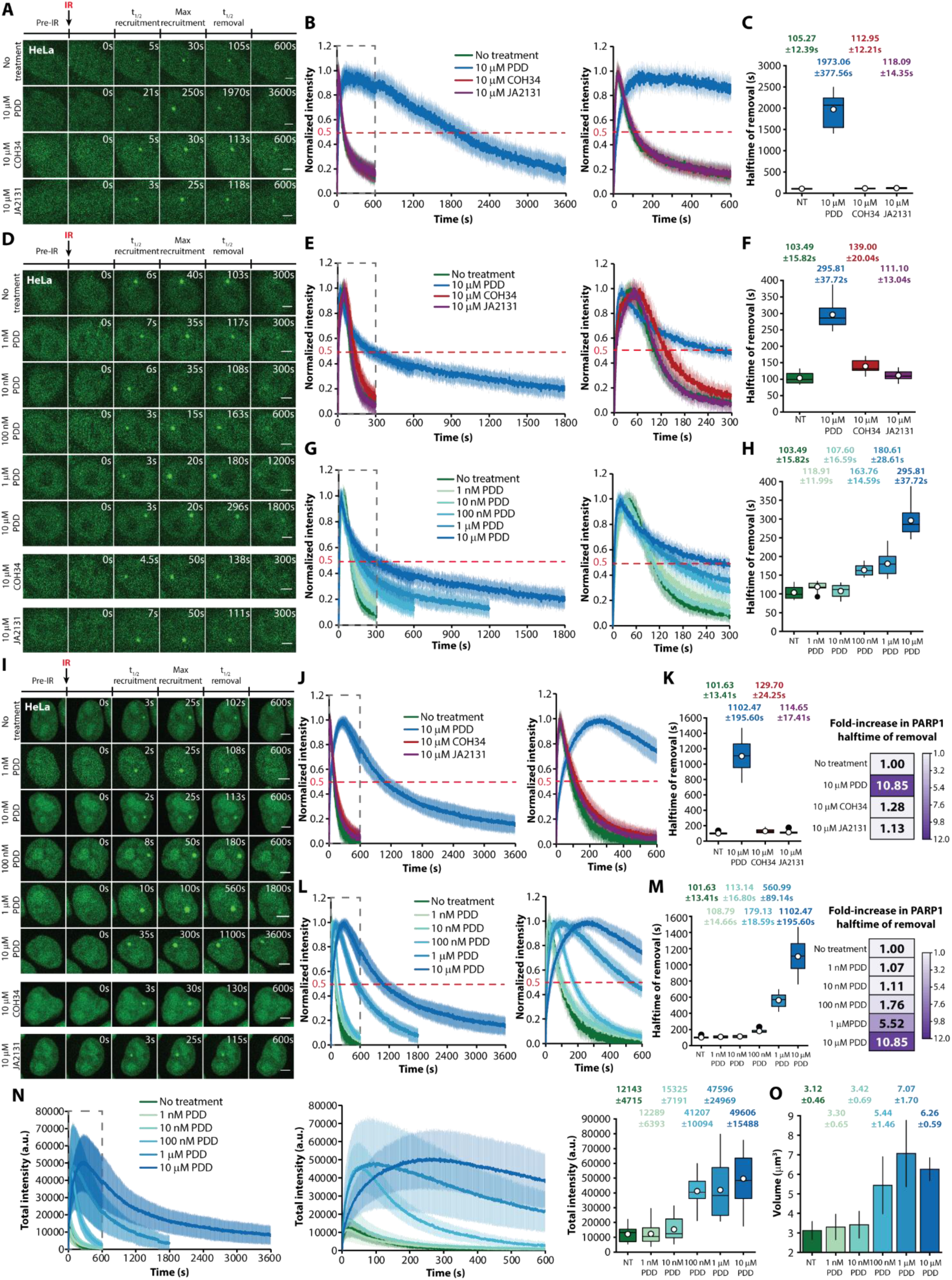
PARG inhibition prolongs PARG, ARH3, and PARP1 residence at DNA damage sites. Time-lapse images, normalized kinetics, and removal half-times of PARG (A-C), ARH3 (D-H), and PARP1 (l-M) in HeLa cells treated with PDD, COH34or JA2131. For PARP1 IR foci, fold-changes in the half-time of removal are also shown, in addition to total intensity kinetics (N) and volume (O). Data are presented as the mean±SD. White dots indicate the mean value. Scale bars: 5 pm. NT, no treatment; IR, UV laser micro-irradiation; a.u., arbitrary units.

Surprisingly, pre-treatment with PDD extended the presence of PARP1 at DNA damage sites, greatly prolonging PARP1 removal half-times (Figures 1I-1K). This unexpected effect was concentration-dependent, emerging at 100 nM and above (Figures 1I, 1L, and 1M). Notably, 10 µM PDD increased the half-time of PARP1 removal from 101.63±13.41 s in untreated cells to 1102.47±195.60 s, corresponding to a pronounced over 10-fold increase (Figures 1I-1M). Importantly, this effect was consistent across HeLa Kyoto (hereafter referred to as HeLa), PC3, and DLD1 cells (Figures S1F-S1J). While such a delay in PARP1 removal was reminiscent of PARPi-induced retention,^23^ a considerable >4-fold increase in the amount of PARP1 recruited at lesions was also observed (Figure 1N), in addition to а >2-fold enlargement of PARP1 foci volume at damage sites (3.12±0.46 μm^3^ vs. 6.26±0.59 μm^3^ in untreated and 10 µM PDD-treated cells, respectively) (Figure 1O). These results contrast a previous report, which suggested that PARGi reduces the amount of PARP1 recruited upon IR.^45^ Our observations prompted us to assess PARP1 turnover at IR sites via fluorescence recovery after photobleaching (FRAP), since delayed removal was associated with slower exchange upon PARPi treatment.^23^ Indeed, PDD concentrations over 100 nM dramatically slowed PARP1 turnover at damage sites, with 10 µM PDD increasing the fluorescence recovery half-time to 43.12±9.17 s, as compared to 4.62±1.32 s in untreated cells (Figures 2A-2F). This effect on PARP1 turnover was preserved across all tested cell lines (Figures S2A-S2D), while PDD had no effect on the fluorescence recovery of freely diffusive PARP1-EGFP molecules (not bound to damaged chromatin) (Figures S2E-S2G). Of note, PDD slowed down the exchange rate of PARP1 to a much greater extent than the most potent PARPi trapper talazoparib, inducing a 9.33-fold increase in the half-time of PARP1-EGFP fluorescence recovery versus a 5.50-fold increase for talazoparib (43.12±9.17 s vs. 25.41±5.17 s, respectively)(Figures 2C-2F).^23^ Given that we did not observe any significant changes in the mobile fraction of PARP1 at IR sites (Figure 2E), our data suggest that in PDD-treated cells PARP1 molecules are robustly recruited to damaged chromatin and, once engaged, remain there for substantial periods of time before dissociating.

**Figure 2.**
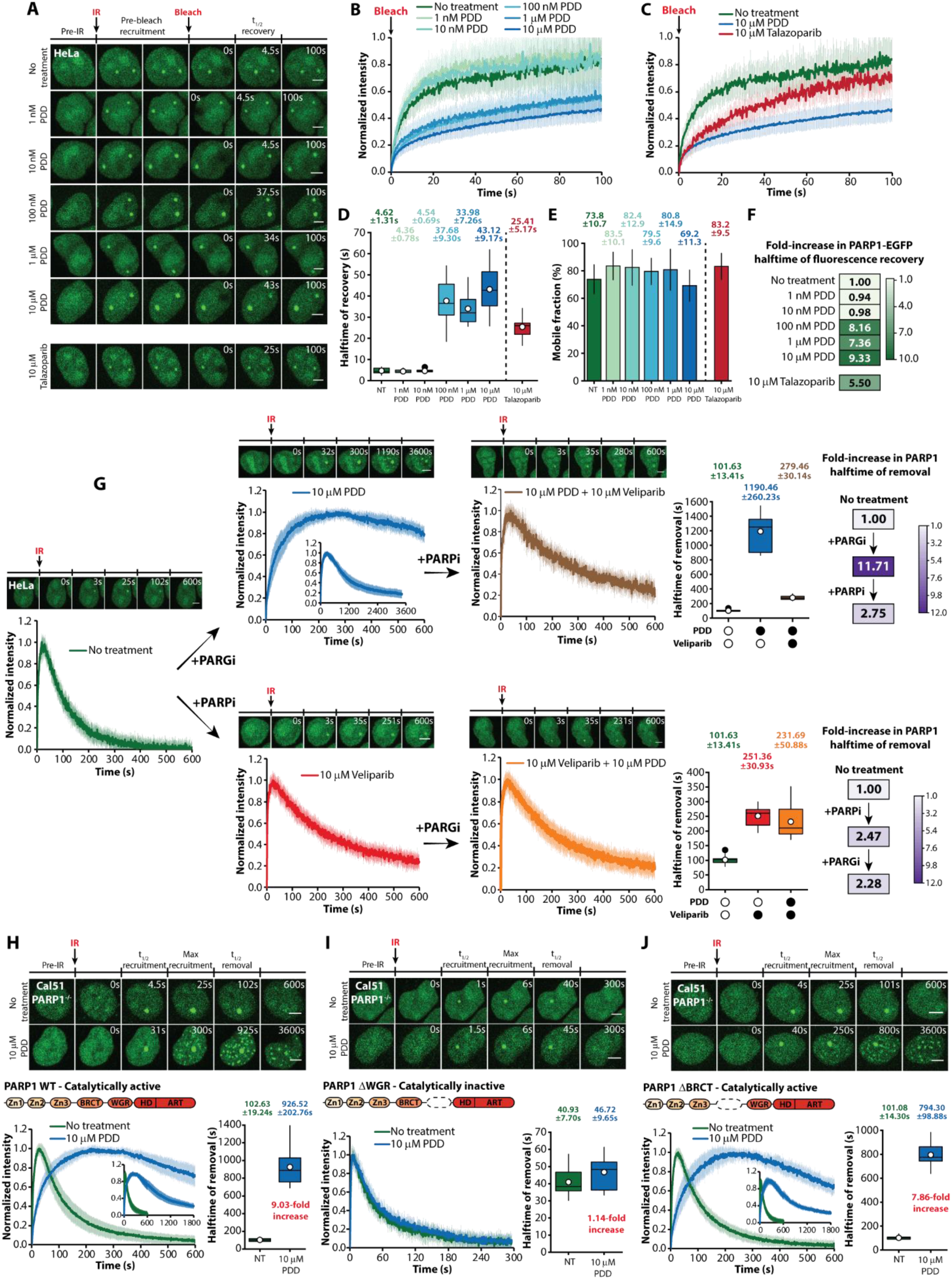
PARG inhibition slows down PARP1 turnover at damaged chromatin sites. Time-lapse images (A), normalized kinetics (B, C), half-times of fluorescence recovery (including fold-increases), and mobile fraction (%) of PARP1 at DNA damage sites (D-F), as determined via fluorescence recovery after photobleaching (FRAP) in HeLa cells treated with PDD and talazoparib. (G) Time-lapse images, normalized kinetics, and PARP1 removal half-times (including fold-increases) in HeLa cells sequentially treated with PDD and veliparib or vice versa. (H-J) Time-lapse images, normalized kinetics, and removal half-times ofWT, AWGR, and ABRCT PARP1 transiently expressed in CalSl PARP1-/-cells treated with PDD. Data are presented as the mean±SD. White dots indicate the mean value. Scale bars: 5 pm. NT, no treatment; IR, UV laser micro-irradiation.

Again, COH34 and JA2131 did not produce any considerable changes in either bulk PARP1 kinetics (Figures S1L-S1S), or PARP1 turnover (Figures S2H and S2I), with minor delays in PARP1 removal noted at 100 µM or when cells were pre-treated with 10 µM COH34 or JA2131 overnight to exclude poor cellular uptake. These results are congruent with PARG, ARH3, and XRCC1 data and call into question the capacity of COH34 and JA2131 to modulate the PARylation–dePARylation cycle in vivo, highlighting PDD as the only bona fide PARG inhibitor of the three. Our observations are also in line with the reported inconsistencies between PDD and these compounds in a recently published PARGi sensitivity CRISPR screen.^39^ Thus, we excluded COH34 and JA2131 from further analyses.

Next, we set out to determine the requirements for the observed PARGi-induced retention of PARP1 at IR foci. Co-treatment with PDD and the non-trapping PARP inhibitors veliparib (Figure 2G) or pamiparib (Figure S2J) completely abolished the PDD-induced changes in PARP1 dynamics, leaving only the effects of the PARPi.^23^ This indicates that the alterations in PARP1 behavior following PDD treatment are due to its PARylation activity. This was further corroborated by experiments in Cal51 *PARP1^-/-^* cells.^49^ The removal of transiently-transfected WT PARP1-EGFP was delayed 9-fold following 10 µM PDD treatment as compared to untreated cells (926.52±202.76 s vs.102.63±19.24 s) (Figure 2H), recapitulating our results obtained in HeLa, PC3, and DLD1 cells. Notably, the PARylation-defective PARP1 ΔWGR mutant failed to exhibit differential dynamics under PARG inhibition (Figure 2I), as opposed to the PARylation-competent PARP1 ΔBRCT mutant, whose removal from damaged chromatin, similarly to that of WT PARP1, was delayed 8-fold upon PDD treatment (Figure 2J). Of note, while PARP1 ΔWGR was recruited to IR-induced lesions, its residence was considerably shorter (t_1/2_removal=40.93±7.70 s), as compared to that of the WT PARP1 in untreated cells (102.63±19.24s), suggesting impaired binding to DNA breaks. Likewise, in untreated cells PARP1 ΔZn2 (t_1/2_removal=63.40±12.83 s) was removed from lesions much more rapidly than WT PARP1 (Figure S2K). However, in stark contrast to PARP1 ΔWGR, PDD delayed ΔZn2 removal almost 8-fold, which aligns with its near-preserved PARylation capacity (Figure S2K).^50^ Finally, the PARP1 E988K mutant, which is only able to generate mono(ADP-ribose) (MAR),^51,52^ exhibited a slight delay in its removal following PDD treatment, greatly attenuated compared to that of its PARylation-competent counterparts (Figure S2L). In summary, our findings reveal an unexpected effect of PARG inhibition in living cells that is entirely dependent on the catalytic activity of PARP1, resulting in markedly prolonged PARP1 residence and decreased turnover at damaged chromatin sites. This dynamic phenotype, while reminiscent of the PARPi-induced trapping, occurred under conditions of unrestrained chromatin PARylation, rather than the lack thereof.

### PARG inhibition captures PARP1 in the vicinity of DNA lesions

Considering the similarities between PARG and PARP1 inhibition in terms of PARP1 dynamics at damage sites, we sought to disentangle these phenomena. Rather serendipitously, we had observed that addition of PDD enhanced the intensity of previously induced IR foci, which had almost dissolved by the time PDD was added. Taking advantage of such a rapid effect, we micro-irradiated cells, timed PDD addition in accordance with the half-time of removal of untreated PARP1 (101.63±13.41 s), and continued monitoring IR foci (Figure 3A). Strikingly, PARG inhibition elicited the enlargement of pre-existing PARP1 foci, some of which became substantially brighter than at any point between IR and PDD addition (Figures 3A and 3B). Like PDD, talazoparib also acted immediately upon addition, but instead of enlarging the IR foci, it halted the time-dependent decrease in PARP1 intensity, in effect ‘flat-lining’ the intensity curve (Figures 3A and 3C). This comparison provided a clear distinction between PARPi-induced trapping and the PARGi-induced phenotype, whereby the former traps PARP1 molecules at a pre-existing lesion, while the latter triggers the renewed accumulation of PARP1, in effect “capturing” more PARP1 molecules within damaged chromatin regions (Figure 3D).

**Figure 3.**
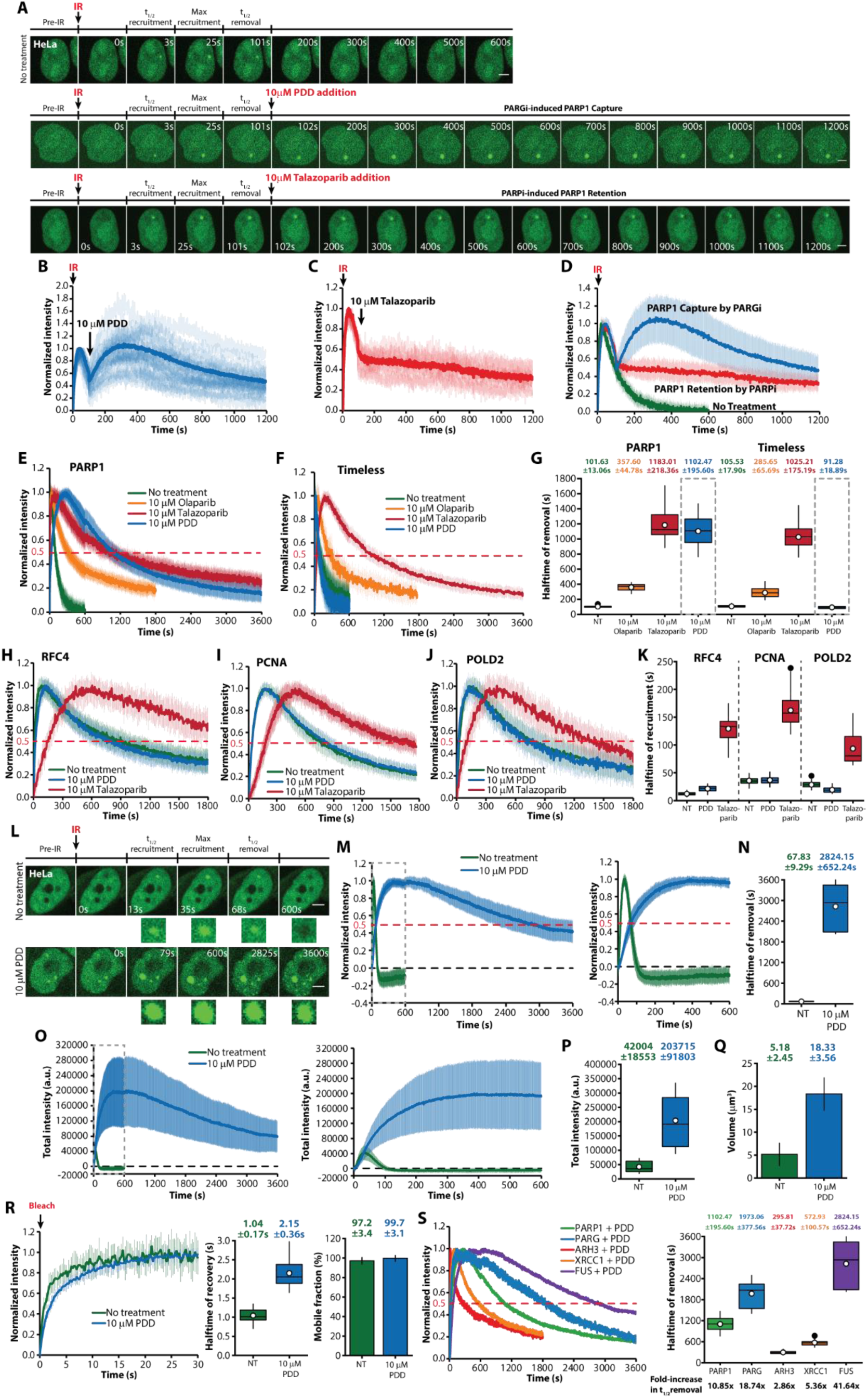
PARG inhibition captures PARP1 at damaged chromatin. (A) Time-lapse images showing the immediate effects of PDD and talazoparib on PARP1 dynamics at pre-existing IR-induced lesions in HeLa cells. (B-D) Normalized kinetics of PARP1 at pre-existing IR lesions following addition of PDD (B), talazoparib (C), and comparison (D) among treatment conditions. (E, F) Normalized kinetics of PARP1 (E) and Timeless (F) at IR sites in HeLa cells pre-treated with olaparib, talazoparib, or PDD. (G) Half-times of removal of PARP1 and Timeless from IR sites. (H-J) Normalized kinetics of REC4 (H), PCNA (I), and POLD2 (J) at IR-induced DNA damage sites in HeLa cells pre-treated with talazoparib or PDD. (K) Jalf-times o recruitment of RFC4, PCNA, and POLD2 to DNA damage sites. (L) Time-lapse images of FUS recruitment and removal from IR-induced damage sites, with and without PDD. (M) Normalized kinetics of FUS at IR lesions, with and without PDD. (N) Half-times of FUS removal from DNA lesions. (O, P) Total intensity kinetics of FUS IR foci, with and without PDD. (Q) Mean volume of FUS foci at IR sites, with and without PDD. (R) (left) Normalized intensity of FUS fluorescence recovery after photobleaching (FRAP), with and without PDD; (middle) Half-times of FUS fluorescence recovery at IR lesions; (right) Mobile fraction of FUS at IR sites. (S) Normalized kinetics and half-times of removal of PARP1, PARG, ARH3, XRCC1, and FUS at lesions after treatment with PDD; fold-increases in removal half-times relative to controls are shown. Data are presented as ment±SD. White dots indicate the mean value. Scale bars: 5 μm, NT, no treatment, IR, UV laser micro-irradiation; a.u., arbitrary units.

To further explore the nature of this “PARP1 capture” effect, we interrogated the dynamics of Timeless and several downstream DNA damage response (DDR) proteins enrolled in DNA synthesis during repair. Timeless is recruited by and directly interacts with lesion-bound PARP1 in a PAR-independent manner, hence, its dynamics provide a clear indicator of *bona fide* PARP1 binding to DNA breaks.^53^ In line with prior evidence, olaparib delayed the removal of Timeless, and this effect was more pronounced under talazoparib treatment, precisely mirroring PARP1 dynamics and confirming that PARP1 and Timeless are co-trapped at DNA lesions (Figures 3E-3G, S3A, and S3B).^23,53^ Notably, PARG inhibition had no effect on the dynamics of Timeless despite the substantial delay in PARP1 removal (Figures 3E-3G) indicating that the PARP1-Timeless interaction at DNA lesions is not altered upon PARGi treatment as compared to the untreated condition. Thus, PARGi-dependent PARP1 capture does not occur at DNA lesions per se, but in their vicinity, in stark contrast to PARPi-induced trapping.

To confirm this, we examined the kinetics of RFC4, subunit of the RFC clamp loader complexes, PCNA, the eukaryotic clamp protein, and POLD2, subunit of DNA polymerase delta. We previously showed that PARPi-induced PARP1 trapping strongly correlates with the delayed recruitment of these proteins to IR sites.^23,48^ Notably, PARG inhibition had no effect on RFC4, PCNA, and POLD2 kinetics, as opposed to talazoparib (Figures 3H-3K and S3C-S3E).^23^ The preserved kinetics of downstream repair factors suggest that the observed excess of PARP1 at IR sites upon PARGi does not directly engage lesions per se, but is rather enriched in the vicinity of damaged chromatin, consistent with the much larger volume of PARP1 IR foci upon PDD treatment (Figure 1O). Taken together, our results point to a role for PARG-mediated dePARylation in preventing the PAR-driven, aberrant accumulation of PARP1 upon DNA damage induction.

### FET family proteins mirror PARP1 behavior under PARG inhibition

The distinct PARP1 capture effect elicited by PDD raises the question as to what factors might contribute to the formation and stabilization of this aberrant DDR assembly. This is particularly relevant for the excess of hyperautomodified PARP1, which carries a substantial negative charge, while persisting within the highly PARylated chromatin environment at damage sites. To address this question, we focused on FET family proteins (FUS, EWSR1, and TAF15), which are well-established mediators of PAR-driven phase separation at sites of DNA damage.^54^ Recently, it was shown that PARP1 and DNA form transient condensates, which mediate the synapsis of broken DNA ends, and these are stabilized by FET family proteins. In particular, FUS was proposed to counteract PAR’s negative charge, thereby stabilizing break-induced, PARylation-dependent PARP1 condensates and fostering their growth.^50^ Notably, FET proteins are rapidly recruited to IR-induced DNA damage sites in a PAR-dependent manner.^48,54,55^ PDD delayed FUS removal from DNA damage sites more than 40-fold (t_1/2_removal=67.83±9.29 s in untreated vs. 2824.15±652.24 s in PDD-treated cells) (Figures 3L-3N) and markedly enhanced FUS recruitment, yielding much larger (5.18±2.45 μm^3^ vs. 18.33±3.56 μm^3^) and brighter (∼5-fold increase in total intensity) IR foci (Figures 3L and 3O-3Q). Of note, while the effects of PDD on FET protein dynamics are consistent with our data for the other PAR-binding proteins, the difference in magnitude was remarkable (Figure 3S), suggesting that FET proteins are particularly responsive to the hyperPARylated environment and could thus facilitate the PARP1 capture effect. Importantly, FUS can bind PAR chains in a multivalent manner through its arginine/glycine-rich (RGG) domains and RNA recognition motif (RRM),^56^ raising the possibility that FUS (and presumably other FET family members) may function as a molecular “sticker” by simultaneous binding to multiple hyperPARylated PARP1 molecules, capturing them in the vicinity of DNA breaks under PARGi treatment. In agreement with this, FUS turnover at damage sites was slowed approximately two-fold upon PARG inhibition (t_1/2_fluorescence recovery=1.04±0.17 s in untreated cells vs. 2.15±0.36 s in PDD-treated cells) (Figures 3R and S3G), suggesting that FUS engages more stably with the excess PAR accumulating at sites of DNA damage. Meanwhile, PARGi did not affect the mobility of non-lesion-bound FUS (Figure S3H). The effects of PDD on the other two FET family members - EWSR1 and TAF15 - were nearly identical (Figures S3I-S3T) and, as expected, veliparib completely abolished FET protein accumulation at damage sites (Figures S3F, S3K, and S3P), effectively negating the consequences of PARGi. In summary, FUS recapitulated PARP1 dynamics upon PARG inhibition, supporting a model in which FET proteins stabilize aberrant PARP1 assemblies.

### PARG inhibition induces PARP1 condensation upon DNA damage

Beyond the dynamic phenotypes described above, our micro-irradiation experiments revealed the formation of secondary, smaller, PARGi-dependent PARP1 foci that formed throughout the nucleus rather than being confined to the vicinity of the IR site (Figure 4A). Previous works reported the formation of such damage-induced foci, showing that these contain hyperPARylated PARP1 and do not co-localize with DNA damage, in addition to describing them as aggregates that maintain constant size and intensity once formed.^45^ In line with this, none of the downstream repair synthesis-associated factors we examined, including PCNA, RFC4, and POLD2 (Figures S3C-S3E), formed such secondary foci following IR. A recently published study also reported a lack of PCNA within PARGi-dependent foci, proposing that these are residual damage assemblies.^39^ However, we observed their formation throughout the entire nucleus following damage induction at a small, localized region of interest, which suggested otherwise. We therefore set out to characterize the nature of these secondary, PARGi-dependent PARP1 foci. Our experiments showed that secondary PARP1 foci are established only by PARylation-proficient PARP1 variants, as observed for WT, ΔBRCT, and ΔZn2 transiently transfected in Cal51 *PARP1*^-/-^ cells (Figures 2H, 2J, and S2K). Meanwhile, PARylation-deficient ΔWGR and E988K PARP1 mutants failed to do so (Figures 2I and S2L). We then followed the formation of secondary foci, which proceeded at a drastically slower rate (t_1/2_formation > 500 sec) in all tested cell lines compared to that of bona fide IR foci under PARGi (Figures 4B and 1L). This difference suggested an underlying mechanism of formation that is distinct from DNA break recognition by PARP1. Next, we subjected HeLa cells to treatments with hydrogen peroxide (H_2_O_2_), methyl methanesulfonate (MMS), and camptothecin (CPT) – compounds that cause diverse PARP1-activating DNA lesions (Figures S4A-S4C).^3,57^ Pre-treatment with PDD resulted in the robust formation of PARP1 foci following addition of each DNA damage-inducing agent, albeit with distinct patterns and timing (Figures 4C and S4A-S4C). Co-treatment with veliparib completely abolished genotoxin-induced foci formation, re-affirming their PAR dependence (Figures S4A-S4C). Considering the non-localized nature of DNA damage induced by these compounds (i.e. throughout the genome), we posited that the observed PARP1 foci are unlikely the consequence of exaggerated PARP1 accumulation at a specific lesion or damaged region, as is the case with the primary focus induced by IR. Similarly to the secondary foci observed after IR, the PARGi-dependent PARP1 foci observed after H_2_O_2_, MMS, and CPT treatment also formed at considerably slower rates than the primary IR-induced PARP1 foci, with multi-fold differences among the three conditions (Figure 4C). Notably, FUS also formed PARGi-dependent secondary foci following IR (Figure 4D), as well as after genotoxin treatment (Figures S4D and S4E), mirroring PARP1 behavior. The kinetics of formation of FUS foci was delayed compared to that of PARP1 foci (t_1/2_formation=931.41±117.61s for FUS foci vs. 702.72±213.44s for PARP1 foci) (Figure 4E), suggesting that FUS is recruited to these damage-induced compartments in a PAR-dependent manner, presumably stabilizing them. To probe the involvement of FUS in a more direct manner, we co-transfected Cal51 *PARP1^-/-^* cells with WT PARP1-EGFP and FUS-mCherry and confirmed their co-localization both at IR sites and in secondary PARGi-dependent foci across the nucleus (Figure S4F). Notably, the PARylation-defective PARP1 E988K mutant greatly suppressed FUS accumulation at IR sites and failed to elicit PARP1 and FUS secondary foci formation under PDD treatment (Figure S4G). The striking resemblance between PARP1 and FET protein behavior in this context provides further support for the notion that FET proteins facilitate PARGi-induced PARP1 condensation upon DNA damage.

**Figure 4.**
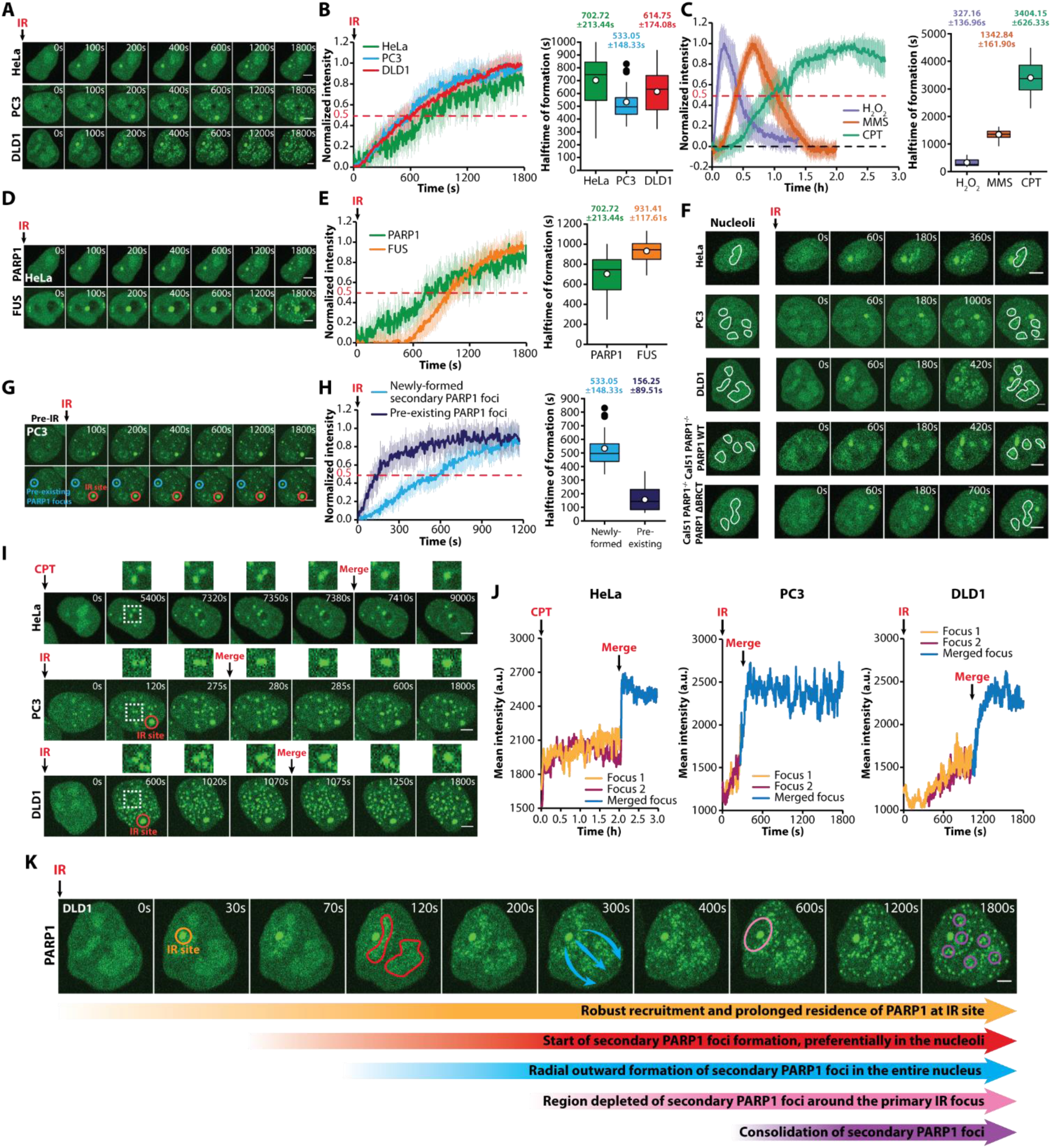
PARG inhibition induces aberrant PARP1 condensation in response to DNA damage. (A) Time-lapse images of PARP1 secondary foci formation after IR in HeLa, PC3, and DLD1 cells treated with PDD. (B) Normalized kinetics and half-times of formation of PARP1 secondary foci after IR in HeLa, PC3, and DLD1 cells treated with PDD. (C) Normalized kinetics and half-times of formation of PARGi-dependent PARP1 foci in HeLa cells treated with 100 pM hydrogen peroxide (H2O2), 0.01% methyl methanesulfonate (MMS), or 10 pM camptothecin (CPT) after lOpM PDD pre-treatment. (D) Time-lapse images of PARP1 and FUS secondary foci following IR in HeLa cells treated with 10pM PDD. (E) Normalized kinetics and half-times of formation of PARP1 and FUS secondary foci after IR in HeLa cells treated with PDD. (F) Time-lapse images showing the nucleolar preference for secondary PARP1 foci formation after IR in HeLa, PC3, DLD1, and Cal51 PARP1-/-expressing WT or ABRCT PARP1 treated with PDD. (G) Time-lapse imaging of pre-existing (before IR) PARP1 foci in PC3 cells treated with PDD, which then grew in response to IR DNA damage induction. (H) Normalized kinetics and half-times of formation of pre-existing versus newly-formed secondary foci following IR in PDD-treated PC3 cells. (I) Time-lapse images showing the coalescence of PARGi-dependent PARP1 condensates after genotoxic treatments in HeLa (10pM camptothecin), PC3 (IR), and DLD1 (IR) cells. (J) Mean intensity tracks of the coalescing foci and the resulting merged focus from (I); a steep, immediate increase in intensity can be observed at the time of fusion. (K) Time-lapse images of PARP1 secondary foci formation after IR in DLD1 cells treated with PDD. Data are presented as mean±SD. White dots indicate the mean value. Scale bars: 5 pm. IR, UV laser micro-irradiation; a.u., arbitrary units.

Interestingly, both the IR-induced secondary PARP1 foci and the genotoxin-induced foci exhibited a tendency to initially form within nucleoli (Figures 4A and 4F). Nucleoli are known to host a higher concentration of PARP1, whilst being DNA-sparse.^58,59^ To exclude the contribution of nucleolar DNA damage, we focused on IR experiments, which allow us to precisely target a region of interest and thus avoid nucleoli. Following damage induction at a non-nucleolar site, we observed PARGi-dependent foci formation preferentially within nucleoli (Figure 4F). While foci proceeded to disperse from nucleoli and spread throughout the nucleus, these observations suggested that foci formation was favored in regions of a higher local PARP1 concentration, which are able to attract, or capture, an increasing share of the available, persistently automodified PARP1 molecules generated at DNA damage sites under PARGi.

During our IR experiments in PC3 cells, we noticed that some PDD-treated cells had formed small, but discernible PARP1 foci even prior to damage induction. We subjected these cells to IR and then followed the fate of such pre-existing foci. Rather than remaining constant in intensity or decreasing in response to the massive accumulation of PARP1 at the IR site, pre-existing PARP1 foci grew (Figure 4G). Furthermore, this growth proceeded at a considerably faster rate than the accumulation of PARP1 within newly-forming secondary foci following IR (156.25±89.51 s vs. 533.05±148.33 s, respectively) (Figure 4H). These observations align well with the nature of biomolecular condensates, where a nucleation barrier prevents rapid accumulation in the case of de novo, unseeded foci, while the kinetics of pre-existing foci are dominated by growth, without an initial, nucleation-attributed lag.^60^ As a further testament to condensate behavior, both the IR-induced secondary foci and the genotoxin-induced foci exhibited a clear tendency to coalesce (Figures 4I and 4J).^61^ Finally, secondary PARP1 foci did not persist in the vicinity of the large primary PARP1 IR focus, suggesting a decreased probability of nearby nucleation due to a reduced local concentration of automodified PARP1 that cannot support the supersaturation necessary for condensate formation (Figure 4K). Considering the above findings together with the well-documented formation of PAR-dependent condensates that abide by the laws of liquid-liquid phase separation (LLPS), we posited that the observed PARGi-dependent foci may, in fact, be of such nature.^62,63^

Taken together, our measurements inform a model whereby a genotoxic trigger of unrestrained PARylation under conditions of PARGi produces large amounts of automodified PARP1, which, upon exceeding a nucleation barrier threshold, forms condensates in cooperation with FET family proteins. Importantly, these condensates do not necessarily form at sites of DNA damage, they grow at much slower rates compared to PARP1 IR foci, coalesce, and, over time, sequester part of the nuclear pool of persistently automodified PARP1. Thus, PARG activity prevents such condensation in response to DNA damage.

### PARP1 condensation is induced by S-phase PARylation and reflects intrinsic PARGi sensitivity

Next, we sought to test and extend this model to a cellular context in the absence of exogenous genotoxic stress. To this end, we followed HeLa, PC3, and DLD1 cells co-expressing PARP1-EGFP and PCNA-mCherry over a 48-h period under PARGi treatment. We took advantage of PCNA localization and foci formation as a well-established indicator of cell cycle progression.^64–66^ In line with the reported requirement of PARG in S phase progression,^37^ PARGi prolonged S phase in PC3 (19.92±5.52 h vs. 10.97±2.79 h in untreated cells) (Figures 5D and 5E) and DLD1 cells (20.03±5.65 vs. 13.37±2.99 h) (Figures 5G and 5H), while having no effect in HeLa cells (7.98±0.87 vs. 8.06±1.00 h) (Figures 5A and 5B). G2 was also prolonged in the former two cell lines, while G1 was unaffected in all three. While practically all HeLa cells that entered S phase under PARG inhibition reached mitosis within 30 h of S-phase entry (Figure 5C), 50.0% and 90.3% of DLD1 and PC3 cells failed to do so, respectively (Figures 5F and 5I). Interestingly, DLD1 and PC3 cells, whose cycle progression was affected by PARG inhibition, formed PARP1 foci in the course of time-lapse imaging, while HeLa cells, which did not exhibit delays in cell cycle progression under PDD treatment, did not form foci (Figures 5O and S5A-S5C). Clonogenic assays confirmed the greater sensitivity of DLD1 and PC3 cells to PDD compared to HeLa cells (Figures 5J-5L). These results suggested that the propensity for PARP1 condensation in the absence of exogenous damage may reflect intrinsic PARGi sensitivity.

**Figure 5.**
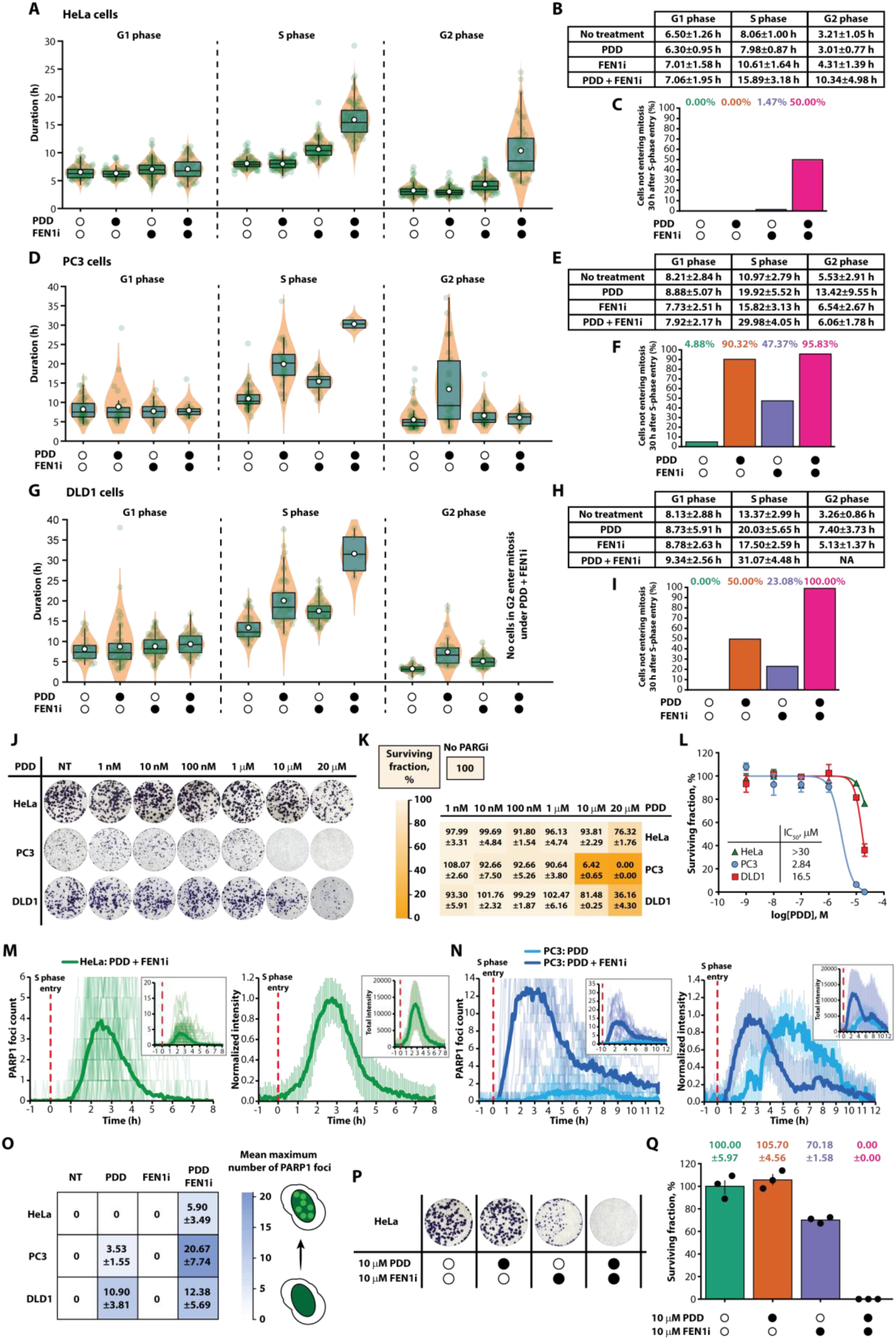
S-phase PARylation triggers PARP1 condensation in a manner that reflects PARGi sensitivity. (A, B) Cell cycle phase durations in PARP1 -EGFP/ mCherry-PCNA-expressing HeLa cells treated with 10 pM PDD, 10 pM FEN1 -IN-1 (FENIi), or both. (C) Fraction of HeLa cells that fail to enter mitosis within 30 h of S phase entry. (D, E) Cell cycle phase durations in PARP1 -EGFP/mCherry-PCNA-expressing PC3 cells treated with 10 pM PDD, 10 pM FEN1 i, or both. (F) Fraction of PC3 cells that fail to enter mitosis within 30 h of S phase entry. (G, H) Cell cycle phase durations in PARP1 -EGFP/mCherry-PCNA-expressing DLD1 cells treated with 10 pM PDD, 10 pM FENIi, or both. (I) Fraction of DLD1 cells that fail to enter mitosis within 30 h of S phase entry. (J-L) Representative images (J) and surviving fractions (K, L) from clonogenic assays of HeLa, PC3, and DLD1 cells subjected to increasing concentrations of PDD. (M) Counts and normalized kinetics of formation of PARP1 foci in HeLa cells co-treated with 10 pM PDD and 10 pM FENIi. Cells are aligned to S phase entry. In the left panel, the count profiles of individual cells are shown in light green, while the mean foci count is shown in dark green. Total intensities of individual foci are also shown in the upper right corner of the right panel. (N) Counts and normalized kinetics of formation of PARP1 foci in PC3 cells treated with 10 pM PDD alone or in combination with 10 pM FEN1 i. Cells are aligned to S phase entry. In the left panel, the count profiles of individual cells are shown in lighter shades, while the mean foci counts are shown with thick lines. Total intensities of individual foci are also shown in the upper right corner of the right panel. (O) Mean maximum number of PARP1 foci in HeLa, PC3, and DLD1 cells treated with 10 pM PDD, 10 pM FENIi, or both. (P, Q) Representative images (P) and surviving fractions (Q) from clonogenic assays of HeLa cells treated with 10 pM PDD, 10 pM FENIi, or both. Data are presented as the mean±SD, except in panels K, L, and 0, where surviving fractions are presented as mean±SEM (n = 3). White dots indicate the mean value. NT, no treatment.

To explore the potential of PARP1 foci as a biomarker of PARGi sensitivity, we first sought to determine the molecular basis underlying their formation. Unequivocally, PARP1 foci formation ensued after S-phase entry in both PC3 and DLD1 cells subjected to PARGi, as confirmed based on foci count and individual foci kinetics (Figures 5N and S5D). As discussed above, we propose that PARGi-dependent PARP1 foci form in response to a sufficient PARylation trigger (e.g. exogenous DNA damage induction). In the course of the cell cycle, pronounced PARylation has been reported to occur during S phase, particularly as a result of PARP1 activation during a backup mechanism for Okazaki fragment maturation.^67,68^ We therefore posited that Okazaki fragment maturation may provide the endogenous PARylation trigger necessary for PARP1 condensation. To probe this, we inhibited FEN1 (FEN1i), which is an essential factor in canonical Okazaki fragment maturation and whose inhibition triggers a several-fold increase in S-phase PARylation due to the greater reliance on PARP1.^67,69^ Combined FEN1i and PARGi treatment drastically increased mean PARP1 foci counts in PC3 cells (Figure 5O), while the requirement for S-phase entry as a prerequisite for their formation remained (Figures 5N and S5B). Importantly, co-inhibiting FEN1 considerably shifted foci kinetics, with foci reaching maximum intensity within ∼2 h of S-phase entry as opposed to ∼5 h under PDD alone, reflecting more rapid foci formation (Figure 5N). This shift in foci dynamics toward S-phase entry was also observed in DLD1 cells under PARG and FEN1 co-inhibition (Figure S5D). This supported the above-proposed model, whereby PARP1 condensation ensues once sufficient levels of automodified PARP1 accumulate. FEN1 inhibition triggers massive PARP1 activation, amplifies the generation of PAR over 10-fold^67^ and thus increases automodified PARP1 levels, leading to faster PARP1 condensation. Importantly, the nucleolar preference for their initial formation as well as a tendency to coalesce were also observed in the context of S-phase-driven PARP1 condensates, confirming the behavior observed following exogenous damage induction (Figure S5E).

As shown above, PARGi alone was unable to induce foci formation, affect cell cycle progression, or elicit cytotoxicity in HeLa cells (Figures S5A, 5B, and 5J-5L). Rather intriguingly, combined FEN1i and PARGi forced foci formation in HeLa cells (Figures 5M, 5O and S5A). These foci formed exclusively upon S-phase entry and exhibited a kinetic profile similar to that of foci in PC3 cells (Figures 5M and 5N). In line with the notion that PARP1 condensation during cell cycle progression may reflect PARGi sensitivity, we observed that co-treatment of HeLa cells with FEN1i drastically prolonged S and G2 phases (Figures 5A and 5B). Further, while PDD alone failed to decrease the number of HeLa cells reaching mitosis within 30 h of S-phase entry, 50.0% of cells failed to reach mitosis within that timeframe under PARG and FEN1 co-inhibition (Figure 5C). These delays in cycle progression were corroborated by pronounced cytotoxicity of the combined treatment (Figures 5P and 5Q), which is in line with published CRISPR screens highlighting the synthetic lethal relationship between FEN1 and PARG.^36,37,39,70^ Taken together, we demonstrate that endogenous PARylation events, particularly those triggered by Okazaki fragment maturation during S phase, are sufficient to trigger PARP1 condensation, and the capacity to induce PARP1 condensates reflects intrinsic PARGi sensitivity. Further, exacerbating S-phase PARP1 activity through FEN1 inhibition can force PARP1 condensation and sensitize cells to PARGi.

### PARG inhibition triggers rapid and reversible nuclear export of PARP1 upon S-phase entry

PARP1 is an abundant, exclusively nuclear protein.^71^ Curiously, during our time-lapse imaging experiments, we noted a reduction in nuclear PARP1 intensity in all cell lines treated with PARGi (Figures 6A, 6B, and S6). The observed reduction was asynchronous (i.e. ensued at different times in different cells over the course of imaging). As it did not occur in untreated cells or cells treated with FEN1i alone, we excluded photobleaching as the underlying cause. Of note, treatment conditions were imaged in parallel, ensuring uniform imaging conditions. Aligning cells based on S-phase entry revealed that the drop in intensity commenced during early DNA replication (within ∼1 h after S-phase entry) (Figures 6B, S6B, and S6D). The reduction amounted to ∼30% of nuclear PARP1 intensity in cells treated with PDD alone, while FEN1 co-inhibition hastened the drop and brought the reduction up to ∼40%. This indicates that the almost instantaneous reduction reflects the extent of S-phase PARylation, which was reported to be more than 10-fold greater in the presence of FEN1i when compared to PARGi alone.^67^ By monitoring cytoplasmic fluorescence, we demonstrated that in HeLa cells, the PARGi-induced drop in nuclear PARP1 coincided with an ∼1.3-fold increase in cytoplasmic PARP1 intensity, which was further enhanced through FEN1 co-inhibition (∼1.7-fold increase) (Figure 6B). Cytoplasmic translocation was previously reported for combined PARGi and MMS treatment, but not for PARGi alone, and attributed to cell death.^45^ However, as shown above, cycle progression and colony formation were virtually unaffected in HeLa cells subjected to PARGi alone. The rapid nature of this translocation, coupled with unperturbed cell cycle progression and preserved viability in HeLa cells, suggested that the observed phenomenon was not associated with cell death. Importantly, this cytoplasmic translocation of PARP1 was a reversible process, with a gradual reduction in cytoplasmic intensity observed in parallel to an increase in nuclear signal noted as cells progressed through S and G2 (Figures 6C and 6D). While HeLa cells treated with PDD alone entered mitosis before nuclear PARP1 would return to its pre-S-phase intensity, those under PARG and FEN1 co-inhibition experienced a considerable recovery of nuclear PARP1 intensity during their prolonged S and G2 phases prior to division.

**Figure 6.**
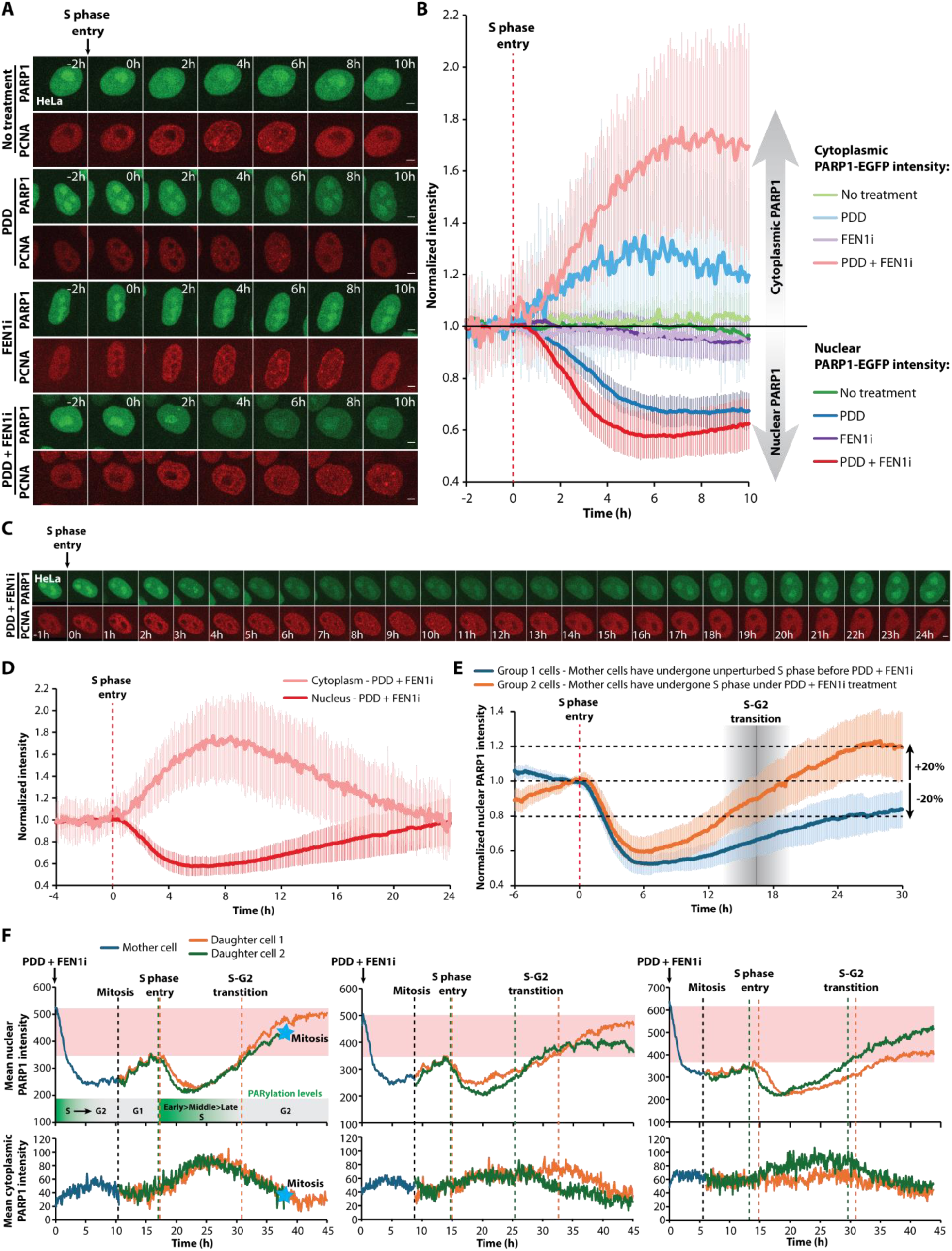
PARG-mediated dePARylation prevents the nuclear export of PARP1 during early S phase. (A) Time-lapse images of PARP1-EGFP/PCNA-mCherry-expressing HeLa cells progressing through S phase without treatment or whilst under treatment with 10 pM PDD, 10 pM FEN1-IN-1 (FENIi), or both. A visible reduction in nuclear PARP1 intensity can be noted under PDD and combined treatment. (B) Cytoplasmic and nuclear PARP1 fluorescence intensity without treatment or whilst under treatment with 10 pM PDD, 10 pM FEN 1 i, or both, normalized to the average intensity of 15 frames before S phase entry. (C) Time-lapse images of HeLa progressing through the cell cycle after S phase entry while under treatment with 10 pM PDD and 10 pM FENIi. Loss and recovery of nuclear PARP1 intensity can be observed over the course of imaging. (D) Normalized fluorescence intensity of nuclear and cytoplasmic PARP1 in HeLa cells treated with 10 pM PDD and 10 pM FENIi. (E) Normalized nuclear PARP1 intensity of Group 1 and Group 2 HeLa cells treated with 10 pM PDD and 10 pM FENIi in the course of cell cycle progression. Group 1 includes cells whose mother cells have undergone unperturbed S phase, prior to PDD+FEN1i addition. Group 2 includes cells whose mother cells have undergone S phase under PDD+FEN1 i treatment. Normalized to the average intensity of 15 frames before S phase entry. (F) Single-cell tracks of mean PARP1 intensity in mother and corresponding daughter cells. Nuclear intensities (upper panels) and cytoplasmic PARP1 intensity (lower panels) are shown. Data are presented as the mean±SD. Scale bars: 5 pm.

In light of the above-described phenomenon, we questioned whether the inability to dePARylate PARP1 would influence PARP1 homeostasis between cell generations. Of the three cell lines tested, HeLa cells provided the ideal, robust model to interrogate this for two reasons: (1) they progressed through S phase even under PARGi+FEN1i treatment, with 50% of cells successfully reaching mitosis during the observed period (Figure 5C); (2) their limited movement made them convenient for image analysis. Thus, we divided HeLa cells co-treated with PARGi and FEN1i into two groups, as follows. Group 1 included cells undergoing their first S phase after the addition of PARGi+FEN1i, meaning that their mother cells had undergone unperturbed S phase, i.e. under no treatment. Group 2 included cells that originated from a mother cell that had already undergone DNA replication under the combined treatment. The goal of this grouping was to assess whether undergoing consecutive S phases under PARGi would result in an additive effect on PARP1 behavior, as each S phase would contribute additional DNA-replication-derived PAR and, thus, automodified PARP1. Indeed, the two groups exhibited distinct nuclear PARP1 dynamics (Figure 6E). Gradual accumulation of nuclear PARP1 was observed during G1 in Group 2 cells, but not in Group 1 cells. Following a pronounced replication-associated drop during early S, both groups experienced recovery of nuclear PARP1 levels, which began approximately halfway into S phase and persisted into G2. In Group 1 cells, nuclear PARP1 levels subsequently recovered to approximately 80% of pre-S-phase levels. Strikingly, Group 2 cells experienced a recovery exceeding pre-S levels by roughly 20% (i.e. these cells exhibited 120% of pre-S-phase PARP1 intensity). A pattern emerged: nuclear PARP1 levels recovered during mid/late S, G1, and G2 phases. Notably, the latter two are characterized by a rather limited PARylation activity,^37,67,68^ while pronounced PARP1 activity has been reported during early S phase.^72^

The increases in nuclear PARP1 levels observed during both growth phases in Group 2 cells, especially during the G2 phase, led us to hypothesize that automodified PARP1 ‘inherited’ from mother cells and re-entering the nucleus could be the source of these additional ∼20% relative to pre-S levels. To probe this more directly, we selected individual mother and daughter cells and plotted both their nuclear and cytoplasmic PARP1 intensities, along with known mitosis entry, S-phase entry, and S/G2 transition (Figure 6F). An immediate drop in nuclear PARP1 was observed in mother cells, which were undergoing S phase at the time of PARGi+FEN1i addition. This suggests that hypermodified PARP1 undergoes rapid nuclear export before it can be efficiently dePARylated. Consequently, these mother cells entered mitosis with considerably lower levels of nuclear PARP1. Importantly, nuclear PARP1 levels prior to mitosis entry closely corresponded to the early G1 levels in daughter cells. This was followed by the above-described gradual increase in G1 and another rapid drop after S-phase entry by the two daughter cells. Only as daughter cells progressed through the greatly extended S and G2 phases did their nuclear PARP1 levels recover back to the mother cell levels measured at the start of imaging. In effect, daughter nuclear PARP1 levels in G2 were approximately two-fold greater than their mother G2 levels, while comparable to the mother cell’s nuclear PARP1 at the start of imaging (Figure 6F). This indicated that substantial nuclear re-entry of PARP1 is taking place during late S/G2. More importantly, the amount of PARP1 that re-enters is greater than the amount exported during the early S phase of the daughter cells. The observed changes in nuclear PARP1 intensity in our single-cell tracks were accompanied by reciprocal dynamics of the cytoplasmic signal (Figure 6F). Notably, recent proteomic profiling did not report a change in PARP1 levels in PDD-treated HeLa cells.^73^ Aligning with our data, another study reported no considerable fluctuations in PARP1 levels through the cell cycle of untreated HeLa cells, excluding cell cycle control as a possible reason for the observed changes in nuclear PARP1.^68^ This strongly suggested that PARP1, which was extruded during the mother cell’s S phase without re-entering by the end of its cell cycle, did re-enter during G1 and the prolonged S/G2 phases of daughter cells.

In summary, our measurements suggested a trans-generational persistence of automodified PARP1, whose nuclear re-entry occurred once sufficient time to dePARylate was available and significant de novo PARylation was not taking place (i.e. during G1, mid/late S, and G2 phases). We demonstrate that, beyond inducing PARP1 condensation, PARG inhibition triggers the nuclear expulsion of persistently automodified PARP1 during S phase, which may leave a considerable fraction of the nuclear PARP1 pool unable to attend to DNA breaks and unligated Okazaki fragments. This reversible phenomenon, in turn, reveals a previously undescribed role of PARG catalytic activity in sustaining the nuclear localization of PARP1 in cycling cells and a potential mechanism of action of PARGi.

## Discussion

### PARP1 is captured at damaged chromatin during unrestrained PARylation

Using live-cell imaging, we demonstrated that PARG inhibition prolongs the overall residence of PARP1 at IR-induced DNA lesions (Figures 1I–1M), while also drastically reducing its turnover (Figures 2A–2F). This unexpected effect, robust across HeLa, PC3, and DLD1 cells (Figures S1F-S1J and S2A-S2D), is, at first glance, reminiscent of PARP inhibitor-induced PARP1 retention.^23^ Further quantification revealed substantial increases in both the total amount of recruited PARP1 and the volume of PARP1 foci at IR sites (Figures 1N and 1O). To validate the hyperPARylated environment at damage sites, we assessed the kinetics of PAR binders and degraders, finding that PARG, XRCC1, and ARH3 dynamics (Figures 1A-1H and S1A-S1D), as well as those of FET family proteins (Figures 3L-3N, S3I-S3M, and S3P-S3R), were considerably prolonged upon PARGi. Although these altered dynamics consistently support the hyperPARylated state of damaged chromatin under PDD, their magnitude varied considerably among proteins (Figure 3S). These differences likely pertain to distinct PAR-binding modules and corresponding PAR topology preferences.^74,75^ Our experiments with two other PARGi compounds - COH34^29^ and JA2131^30^, failed to recapitulate the above-described effects of PDD (Figures 1A-1F, 1I-1K, S1, S2H, and S2I), questioning their capacity to suppress PARG activity in living cells, which is in line with recently reported discrepancies in а CRISPR sensitivity screen.^39^ Finally, through experiments with PARPi co-treatment and PARP1 mutants, we confirmed that the observed alterations in PARP1 behavior under PARGi treatment are entirely dependent on its catalytic activity.

Intriguingly, when PDD addition was timed based on the half-time of removal of PARP1 but without the induction of additional DNA damage, we could precisely follow a rapid second wave of PARP1 accumulation at damage sites, with some foci becoming substantially brighter than at any point before PDD was added (Figure 3B). These data indicate that PARGi induced the growth of pre-existing IR-induced PARP1 foci, suggestive of the capacity of such hyperPARylated DDR assemblies to instantaneously sequester an increasing share of PARP1 molecules. In contrast, talazoparib induced the immediate trapping of PARP1 molecules at damage sites, with no subsequent increase in foci size or intensity (Figure 3C). Taken together, these observations demonstrate that the PARP1 capture effect is mechanistically distinct from the PARPi-mediated retention of PARP1 at DNA breaks (Figure 3D).^23^ This is in line with the distinct genetic vulnerabilities described for PARPi versus PARGi.^39^

To further interrogate the PARP1 capture effect, we used Timeless as a read-out for direct PARP1 binding to DNA breaks in living cells since it interacts with lesion-bound PARP1 in a PAR-independent manner and is co-trapped with PARP1 upon PARPi treatment.^53^ Our findings indicate that, although PARP1 resides at damaged chromatin sites for markedly prolonged periods upon PARGi treatment, PARP1 binds directly to DNA breaks for much shorter periods of time (i.e., while Timeless is also present) that are equal to those observed in untreated cells (Figures 3E-3G). In other words, the prolonged residence of PARP1 at damaged chromatin following PARG inhibition does not reflect prolonged binding to DNA breaks, but rather an enrichment of PARP1 in their vicinity. This was further confirmed by the unaltered dynamics of the downstream repair synthesis-associated factors RFC4, PCNA, and POLD2 (Figures 3H-3K).^23^ The kinetic behavior of the ΔZn2 PARP1 variant provides a final line of evidence that PARP1 capture occurs in their vicinity rather than directly at DNA breaks. Remarkably, although ΔZn2 displays a prominent defect in DNA lesion binding compared to WT PARP1, its retained PARylation activity was sufficient to delay its removal approximately 8-fold upon PDD addition (Figures 2H and S2K). These data suggest that, in contrast to PARPi-induced PARP1 trapping, which physically blocks access to DNA breaks, PARP1 capture following PARG inhibition does not occlude DNA lesions, allowing normal access of downstream DDR proteins.

Under PARGi treatment, the IR-induced foci inevitably host higher levels of hyperautomodified PARP1.^76^ Their greater size and intensity thus entail the presence of one or more factors that can neutralize the clustered negative charge of accumulated PAR chains in order to maintain PARP1 molecules in close proximity. FET proteins are ideal candidates, with FUS recently shown to stabilize PARP1 condensates by accommodating negative PAR chains through its positively charged RBD domain.^50^ Indeed, FET proteins mirrored the dynamic changes observed for PARP1 under PARG inhibition, exhibiting markedly delayed kinetics whilst accumulating at higher amounts and within a much larger volume (Figures 3L-3Q and S3F-S3T). Although we cannot exclude contributions from other PAR-binding proteins in living cells, the resemblance to PARP1 behavior, combined with the multivalent PAR-binding capacity of FET proteins,^56^ favors a model in which FET proteins capture hyperPARylated PARP1 molecules as they dissociate from DNA breaks, thereby forming aberrant long-lived PARP1 assemblies under PARG inhibition. In summary, we define PARP1 capture as a process that unfolds in the context of hyperPARylated chromatin and is entirely dependent on PARP1 catalytic activity, occurring in the vicinity of IR-induced lesions rather than directly at DNA breaks, as is the case with PARPi-induced PARP1 trapping.

### PARG prevents aberrant PARP1 condensation

Corroborating such a proposition, our micro-irradiation experiments revealed the formation of numerous PARGi-dependent PARP1 foci throughout the nucleus following diverse PARP1-activating insults, including IR, H_2_O_2_, MMS, and CPT (Figures 4A-4C and S4A-S4C). While these PARGi-dependent PARP1 foci were recently described, evidence regarding their molecular basis remains inconclusive.^39^ Herein, we provide extensive in vivo evidence of their behavior as PAR-driven biomolecular condensates. To begin with, we demonstrate that, while devoid of downstream repair synthesis-associated factors (PCNA, RFC4, and POLD2), these were also enriched for, and, thus, potentially stabilized by FUS (Figures 4D, S4D, and S4E). Dumoulin et al. also reported that these foci are devoid of PCNA, while enriched for PAR-dependent BER factors XRCC1, LIG3, and POLB, proposing that these may represent residual DNA damage assemblies.^39^ The enrichment of FUS within these foci further motivated us to interrogate their molecular basis as condensates (Figures 4D and S4F). Notably, the growth of PARGi-dependent foci proceeded at a much slower rate than that of PARP1 foci at IR sites (Figure 4B). This can be explained by the fact that the IR-induced DNA damage sites are the regions where automodified PARP1 is generated and rapidly condensed, whereafter, as it slowly dissociates from the primary focus (Figures 1I-1O and Figures 2A-2F), automodified PARP1 proceeds to seed secondary foci elsewhere in the nucleus - at a much slower rate (Figures 4A and 4B). What is more, PARGi-dependent foci exhibited a clear tendency to coalesce, which is a feature of LLPS-driven condensates (Figures 4I and 4J).^61^ In line with the notion of supersaturation being a prerequisite for biomolecular condensate formation,^77^ we demonstrate the propensity of these foci to originate from the nucleoli (Figure 4F), which are PARP1-enriched, and, in the event of persistent nuclear PARylation, represent a likely site to first reach supersaturation of hyperautomodified PARP1 (other than the site of IR). These features provide an argument against the previously proposed nature of PARGi-dependent foci as protein aggregates.^45^ In addition to the convincing in vitro experiments demonstrating that FUS can buffer the negative charge within PAR-rich assemblies,^50^ recent work described PARGi-driven FET condensation at replication forks.^78^ Thus, the capture phenomenon is not exclusive to sites of IR-induced DNA damage, but occurs within regions where the local concentration of hyperautomodified PARP1 exceeds a threshold, be it the nucleoli or the vicinity of an extensively damaged region following IR. As PARP1 capture does not occur in untreated conditions, we posit that this is a property of persistently automodified PARP1 molecules, considering that PARP1 is among the major recipients of damage-induced PARylation.^76^ In other words, PARGi-induced PARP1 capture is an aberrant biomolecular condensation event.

Notably, all of the above-described condensate features were confirmed for foci that arise both as a result of exogenous and endogenous PARylation events. PARGi-dependent condensate formation in the absence of exogenous damage induction was recently proposed to arise as a consequence of endogenous PAR-generating repair events.^39^ Herein, based on extended time-lapse imaging across three cancer cell lines, we demonstrate that, in the absence of exogenous DNA damage, PARGi-induced PARP1 condensation occurs exclusively after cell entry into S phase (Figures 5M, 5N, and S5). Building on this observation, we provide evidence demonstrating that S-phase PARylation constitutes the major endogenous PARylation trigger driving PARP1 condensation under PARGi. By co-inhibiting FEN1, which greatly enhanced condensate formation, we identified PARP1-dependent backup Okazaki fragment maturation as the major source of hyperPARylated PARP1 for condensation (Figures 5M-5O).^67,69^ Тhe observed DNA replication-driven PARP1 condensates were transient, dynamic assemblies, even under the highest levels of PARylation generated upon PARGi and FEN1i co-treatment (Figures 5M and 5N).^67^ What is more, we demonstrate that the propensity of cells to form such condensates in response to PARGi treatment tightly corresponds to their intrinsic sensitivity to PARGi, as reflected by delays in cell cycle progression and survival (Figures 5A-5L and 5O). That is, only cells exhibiting PARGi-induced replication stress (as reflected by prolonged S phase duration) formed PARP1 condensates under treatment with PDD alone, which was the case for PC3 and DLD1 cells (Figures 5D-5I and 5O). Meanwhile, HeLa cell cycle progression was unaffected by PARGi alone, and this was paralleled by an absence of PARP1 condensates (Figures 5A-5C and 5O). Notably, through FEN1i co-treatment, which has been shown to amplify S-phase PARylation over 10-fold,^67^ we were able to force PARP1 condensation in HeLa cells (Figures 5M, 5O, and S5A). This indicates that PARP1 condensation reflects an insufficient cellular dePARylation capacity and, importantly, accompanies the sensitization of an otherwise resistant cell line to PARGi (Figures 5P and 5Q). These results support a role for PARP1 condensation as a biomarker of PARGi sensitivity, rather than a universal consequence of PARG inhibition. Considering that the formation of PARP1 condensates requires PARG inhibition, we propose that they are aberrant assemblies fueled by nuclear PARylation events in the absence of timely reversal, originating within regions where a supersaturation threshold is first breached, rather than exclusively forming at sites of damage. Thus, PARP1 condensate formation signifies insufficient dePARylation capacity that would otherwise prevent it. Perhaps even more intriguingly, the current findings suggest that PARGi may act, at least in part, through the induction of aberrant biomolecular condensate formation. While dysregulated phase separation is now a well-established molecular feature of disease,^79^ to our knowledge, there is limited evidence of anticancer drugs that elicit aberrant phase separation. Taken together, our observations point toward a role for PARG in the prevention of aberrant PARP1 condensation, which is not a shared feature of cancer cell lines, but is indicative of a given line’s dePARylation capacity and, consequently, its sensitivity to PARG inhibition.

### PARylation-driven nuclear extrusion of PARP1

Herein, we report the rapid nuclear extrusion of PARP1 as another consequence of PARG inhibition that occurs as cells enter S phase (Figures 6A, 6B, and S6). Importantly, loss of nuclear PARP1 was particularly robust and observed not only in PC3 and DLD1, but also in HeLa cells, whose cycle progression and viability were virtually unaffected by PDD treatment alone (Figures 5A-5C and 5J-5L). We demonstrated that PARP1 is extruded into the cytoplasm in the absence of cell death or even minor cell cycle delays in HeLa, which points toward the persistent automodification of PARP1 acting as a determinant of its subcellular localization (Figure 6B). What is more, nuclear-extruded PARP1 proceeded to re-enter nuclei during G1, mid/late S, and the prolonged G2 phase, as could be observed in cells subjected to PARG and FEN1 co-inhibition, in which the amount of nuclear PARP1 loss approached 40% (Figures 6B-6D). A striking increase of up to 120% of pre-S levels was observed in some cells by the time they experienced prolonged G2 (Figure 6E). Taking advantage of recently published proteomic analysis of the response to PDD, which shows no effect on PARP1 levels in HeLa cells, we surmise that the observed increase in nuclear PARP1 during G2 did not reflect a change in cellular PARP1 levels.^73^ Instead, by following mother and daughter cells as they progressed through the cell cycle under conditions of persistent PARylation, we demonstrate that automodified PARP1 can be inherited, with the growth phases providing windows of opportunity for PARG and/or ARH3-mediated dePARylation and the nuclear re-entry of PARP1 (Figure 6F). Such dePARylation may also occur in the cytoplasm, where two catalytically active isoforms of PARG localize.^80^ Notably, while PARP1 condensation is not a universal consequence of PARG inhibition (Figure 5O), the loss of nuclear PARP1 is, regardless of sensitivity (Figures 6A, 6B, and S6). PARG inhibition was recently proposed to decrease repair capacity as a consequence of condensate formation.^39^ Our findings suggest that PARylation-driven, reversible extrusion of a substantial fraction of nuclear PARP1 molecules, which become unavailable to engage DNA breaks and Okazaki fragments, may represent an additional mechanism through which PARG inhibition elicits cytotoxicity. Beyond the mechanistic study of PARGi, these observations add to a limited body of evidence of a direct effect of PARylation on subcellular localization.^81,82^ Rather intriguingly, while PARP1 is not under cell cycle control (i.e. neither in levels, nor in subcellular localization),^68^ failure to dePARylate may indeed place PARP1 localization under cell cycle regulation that is dependent on its PARylation status. The exact mechanism and significance of this loss of nuclear PARP1 remain unclear, yet our results provide evidence for a hyperPARylation-driven nuclear export mechanism that is particularly intriguing and warrants further study.

In conclusion, we demonstrate that PARG activity prevents the aberrant condensation of PARP1 in response to both exogenous genotoxins as well as endogenous triggers of PARylation. We narrow down S-phase-associated PARylation, triggered during the PARP1-dependent backup Okazaki fragment maturation pathway, as the sufficient endogenous source of automodified PARP1 to elicit such condensation upon PARGi. We show that the capacity of PARG inhibition to elicit PARP1 condensation directly reflects intrinsic PARGi sensitivity, and, by aggravating endogenous PARP1 activity through FEN1 co-inhibition, we demonstrate that condensate formation parallels PARGi sensitization and may serve as a bio-marker for PARGi cytotoxicity. Intriguingly, our results demonstrate that insufficient dePARylation reversibly forces PARP1 out of the nucleus during DNA replication, revealing an important means through which PARGi may impede PARP1-driven DNA repair and Okazaki fragment maturation. Ultimately, identifying PARP1 condensation as a robust indicator of PARG inhibition and Okazaki fragment maturation as a targetable underlying process could facilitate the mechanism-based clinical implementation of PARGi.

### Limitations of the study

Our results elucidate the basis of PARGi-induced PARP1 condensation, that is, a sufficient PARylation trigger coupled with compromised dePARylation capacity. While we show that the propensity to form such condensates reflects PARGi sensitivity, we cannot conclusively point to PARP1 condensates as drivers of PARGi cytotoxicity. As previously suggested, biomolecular condensates may form as epiphenomena that are simply the result of a high concentration of condensate components at a given location.^83^ Thus, it is possible that these condensates do not elicit cytotoxicity in and of themselves, but instead reflect insufficient dePARylation capacity. Condensate formation occurs in parallel to the rapid nuclear export of a considerable fraction of PARP1. The exact molecular basis of this export, particularly in light of its reversible nature, warrants further study. In addition, whether PARP1 is the only protein subject to such hyperPARylation-driven export remains to be addressed.

## Acknowledgements

The research leading to these results was funded by the Second Swiss Contribution under the program “Promotion of Young Scientists in Central and Eastern Europe (PROMYS)” (grant no. IZPYZ0_228842 to R.A.). P.-B.K. acknowledges funding from Project No. BG16RFPR002-1.008-0001 BioMedRTC. This research was co-funded by the European Union through the European Regional Development Fund under the RIDST Programme 2021–2027 and the Sofia Euro-Bioimaging node of NRIR-MON (D01-104). The authors gratefully acknowledge the kind assistance of the Bulgarian Advanced Light Microscopy Node of the Euro-BioImaging Consortium, based at the Institute of Molecular Biology, BAS. We are grateful to Nagaraja Chappidi (MPI-CBG) and Simon Alberti (TU Dresden) for kindly sharing PARP1 expression plasmids.

## Experimental model

Stable HeLa Kyoto (RRID: CVCL_1922, sex: Female, Cervical cancer), PC3 (RRID: CVCL_0035, sex: Male, Prostate cancer) and DLD1 (RRID: CVCL_0248, sex: Male, Colorectal cancer) cell lines expressing EGFP-/mCherry-fused proteins of interest from bacterial artificial chromosomes (BACs) as well as Cal51 cells (RRID:CVCL 1110, sex: Female, Breast cancer) with bi-allelic deletions of PARP1 (Cal51 *PARP1^-/-^*) were used for imaging.^46,48,49^ HeLa and Cal51 *PARP1^-/-^* cells were cultured in Dulbecco’s Modified Eagle Medium (DMEM), while PC3 and DLD1 cells were cultured in RPMI1640 medium, both supplemented with 10% fetal bovine serum (FBS), 100 units/ml penicillin, and 100 μg/mL streptomycin. Cells were seeded in 35-mm MatTek glass-bottom dishes (Cat# P35G-1.5-14-C) or 4-chamber 35-mm Cellvis glass-bottom dishes (Cat# D35C4-20-1.5-N) at least 48 h before imaging. Cal51 *PARP1^-/-^* cells were transfected using Effectene Transfection Reagent (QIAGEN) on the following day and imaged 24-48 h after transfection. Transiently transfected Cal51 cells with comparable expression levels to those in BAC-tagged lines, based on fluorescence intensity, were selected for IR and imaging. For HeLa and Cal51 cells, medium was changed to FluoroBrite DMEM, supplemented with 10% FBS, 100 units/mL penicillin, 100 μg/mL streptomycin, and 2mM GlutaMAX, prior to imaging. Meanwhile, PC3 and DLD1 cells were imaged in the same medium they were cultured in (RPMI1640). All inhibitors were dissolved in DMSO, with the final DMSO concentration in the cell culture medium maintained below 0.3% throughout imaging.

## Method details

### Image acquisition

Prior to imaging, petri dishes mounted on the microscope were left to thermally equilibrate for at least 30 min under optimal growth conditions at 37°C, 5% CO_2_, and 90% relative humidity, which were then maintained throughout the imaging session. For micro-irradiation and FRAP experiments, live-cell imaging was performed on an Andor Revolution spinning-disk confocal system with a Nikon Eclipse Ti-E inverted microscope equipped with the Nikon Perfect Focus System (PFS). Image acquisition was performed using a Nikon CFI Plan Apo VC 60x (NA 1.2) water immersion objective and a high-sensitivity iXon897 Electron Multiplying Charge-Coupled Device (EMCCD) camera. The pixel size for this setup was 0.23μm. For micro-irradiation and FRAP experiments, cells expressing EGFP-tagged PARP1, ARH3, XRCC1, and PARG, as proteins with rapid recruitment and removal kinetics, were imaged in a single Z plane. All other proteins were imaged in three Z planes with a step size of 0.5 μm, which were then combined via maximum intensity projection for subsequent image analyses. Between-frame time intervals for micro-irradiation experiments varied 0.25-10 s depending on the type of experiment and the protein of interest.

For extended time-lapse imaging, including that of genotoxic compound treatment and cell cycle progression, we used the Andor Dragonfly 505 dual lens spinning-disk confocal system owing to its wider field of view and higher resolution. This system is equipped with a Nikon Eclipse Ti2 inverted microscope and a Nikon Perfect Focus System (PFS). Image acquisition was performed using a Nikon Apo 60x (NA 1.4) oil immersion objective, and an iXon888 EMCCD camera. The pixel size for this setup was 0.2 μm. Acquisition was performed in 3-7 Z planes with a step size of 0.5 μm, followed by maximum intensity projection for subsequent image analyses. For time-lapses following cell cycle progression in PARP1-EGFP/PCNA-mCherry-expressing HeLa, DLD1, and PC3 cells, images were acquired at 5 min intervals over a ∼48 h period. For these acquisitions, cells were seeded in 4-chamber 35 mm Cellvis glass-bottom Petri dishes (D35C4-20-1.5-N) and different treatment conditions (in different chambers) were imaged in parallel. Camptothecin was added at 10 μM (in DMSO), MMS at 0.01% (in ddH_2_O), and H_2_O_2_ at 100 μM (in ddH_2_O). For all extended imaging experiments, PARGi (PDD00017273) and FEN1i (FEN1-IN-1) were used at 10 μM.

### UV laser micro-irradiation and photobleaching

UV laser micro-irradiation and photobleaching were performed as previously described.^23,84^ To induce complex DNA lesions, we use Andor MicroPoint - a 365 nm dye laser pumped by a 337-nm nitrogen laser emitting 3.5 ns pulses with 150 μJ peak energy. Fluorescence recovery after photobleaching (FRAP) experiments were performed to determine protein exchange at micro-irradiation-induced lesions and unperturbed nuclear regions. To this end, an Andor FRAPPA photobleaching and photoactivation module was employed. A 488 nm laser with 50 mW nominal power attenuated to 30% was used for photobleaching, with 20 repeats and 100 ms dwell time. Bleached ROIs had a circular shape, with a 10-pixel diameter, equivalent to 2.3 μm. To assess exchange at lesions, we micro-irradiated simultaneously two regions in a single cell, one of which was consequently subjected to photobleaching, while the other was used for normalization in light of the fast dissociation kinetics of PARP1 and FUS, as previously described.^23,84^ Bleaching was timed to the maximal accumulation of the protein of interest at the IR site.

### Clonogenic assay

Clonogenic assays were performed as previously described.^23,85^ In brief, HeLa and DLD1 cells were seeded in 6-well plates at 400 cells per well. PC3 cells were seeded at 600 cells/well. PDD00017273 treatment and/or FEN1-IN-1 co-treatment commenced 48 h after seeding, and inhibitor-containing medium was replenished every 4 days. After two weeks, cells were fixed in 10% formalin in phosphate-buffered saline (PBS) for 30 min, then stained with 0.1% Crystal violet dissolved in ddH_2_O for 30 min. IC_50_ values for cell viability were estimated, as previously described, a log[PARGi concentration] versus normalized response model with a variable slope was fitted in GraphPad Prism to pre-calculated surviving fractions.^23,85^

### Quantification and statistical analysis

We analyzed repair protein kinetics at sites of micro-irradiation in CellTool^86^ as previously described.^23,84^ In brief, the mean fluorescence intensity (MFI) of the micro-irradiated region is measured and from it the MFI of a shell-like region immediately surrounding it is subtracted in order to compensate for photobleaching. To obtain the total fluorescence intensity within the damaged region, the mean intensity is multiplied by the area of the micro-irradiated ROI. Consecutive Reactions Chain (CRC) mathematical modelling is then applied to determine the precise half-times of recruitment and removal at the single-cell level, as previously described.^48,84^ IR foci volumes (for PARP1 and FET proteins) were calculated based on the radius of the ROI used for protein kinetics measurements; for the sake of simplicity, IR foci were approximated to ideal spheres. The exchange of fluorescent proteins at damage sites or unperturbed nuclear regions was also analyzed as previously described.^23,84^ A double exponential FRAP equation for a circular ROI was employed to determine fluorescence recovery half-times.^87^ Endogenous foci counts were quantified using Trackmate.^88^ Endogenous foci kinetics were obtained using CellTool, as described above for IR-induced foci. Mean nuclear and cytoplasmic intensities were measured using respective ROIs (nuclear and cytoplasmic) in CellTool.

### Software

CellTool, a freely available image analysis software previously developed by our group, was used for all processing tasks, including intensity-based segmentation and signal tracking as well as mathematical modelling.^86^

---

**Figure SI.**
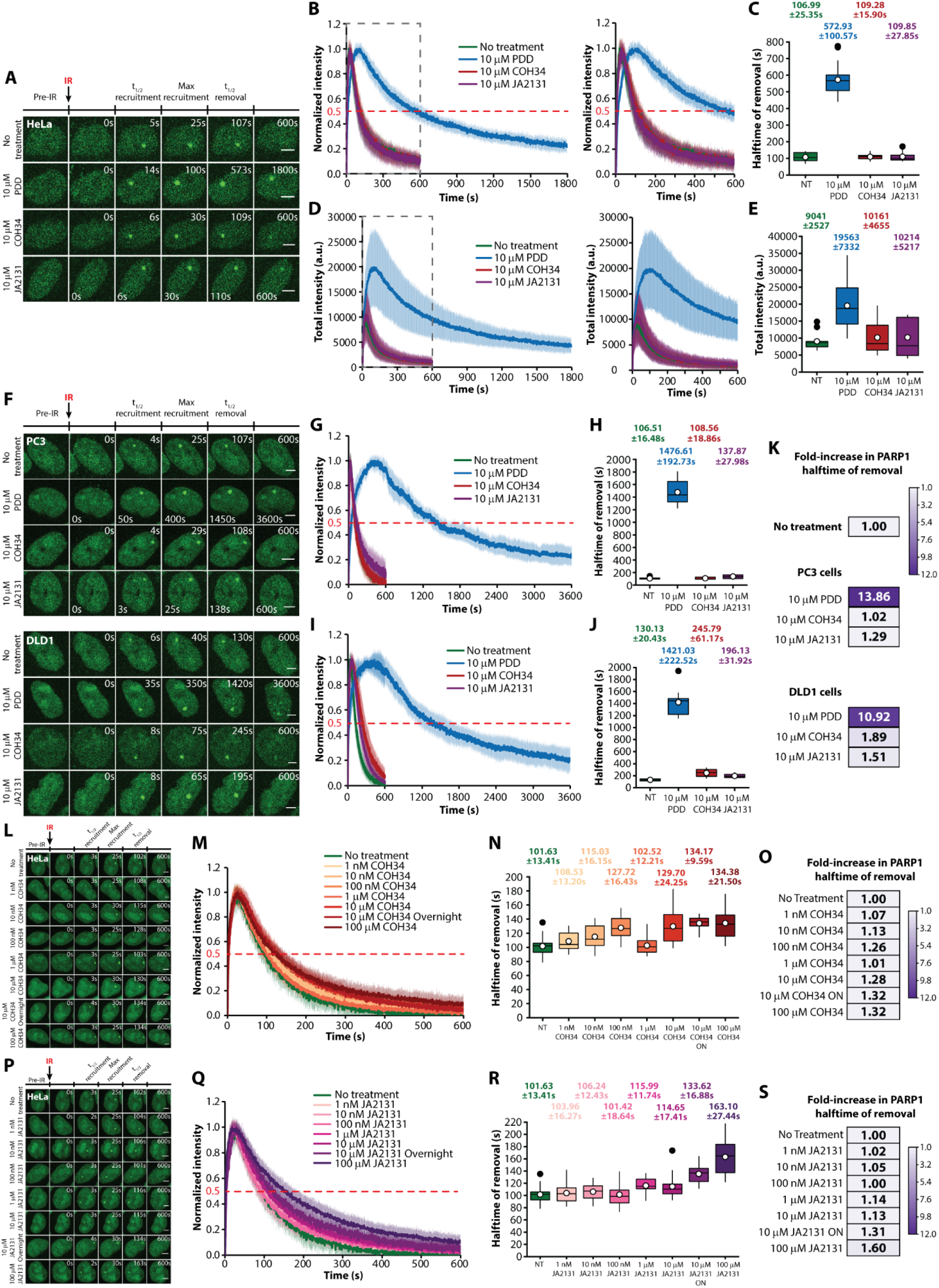
PARG inhibition delays the removal of XRCC1 and PARP1 from DNA damage sites. (A E) Time-lapse images, normalized kinetics, half-times of removal, and total intensity kinetics of XRCC1 IR foci in HeLa cells treated with PDD, COH34, or JA2131. (F) Time-lapse images of PARP1 IR foci in PC3 and DLD1 cells treated with PDD, COH34, or JA2131. (G, H) and (I, J) Normalized kinetics and half-times of removal of PARP **1** in PC3 and DLD1 cells, respectively, treated with PDD, COH34, orJA2131. (K) Fold-changes in RARP1 removal half-times upon PARGi in PC3 and DLD1 cells as compared to the untreated condition. (L-O) and (P-S) Time-lapse images, normalized kinetics, and PARP1 removal half-times (including fold-increases) in HeLa cells treated with a range of concentrations of COH34 and JA2131, respectively. Data are presented as the mean±SD. White dots indicate the mean value. Scale bars; 5 pm. NT, no treatment; IR, UV laser micro-irradiation; ON, overnight; a.u., arbitrary units.

**Figure S2.**
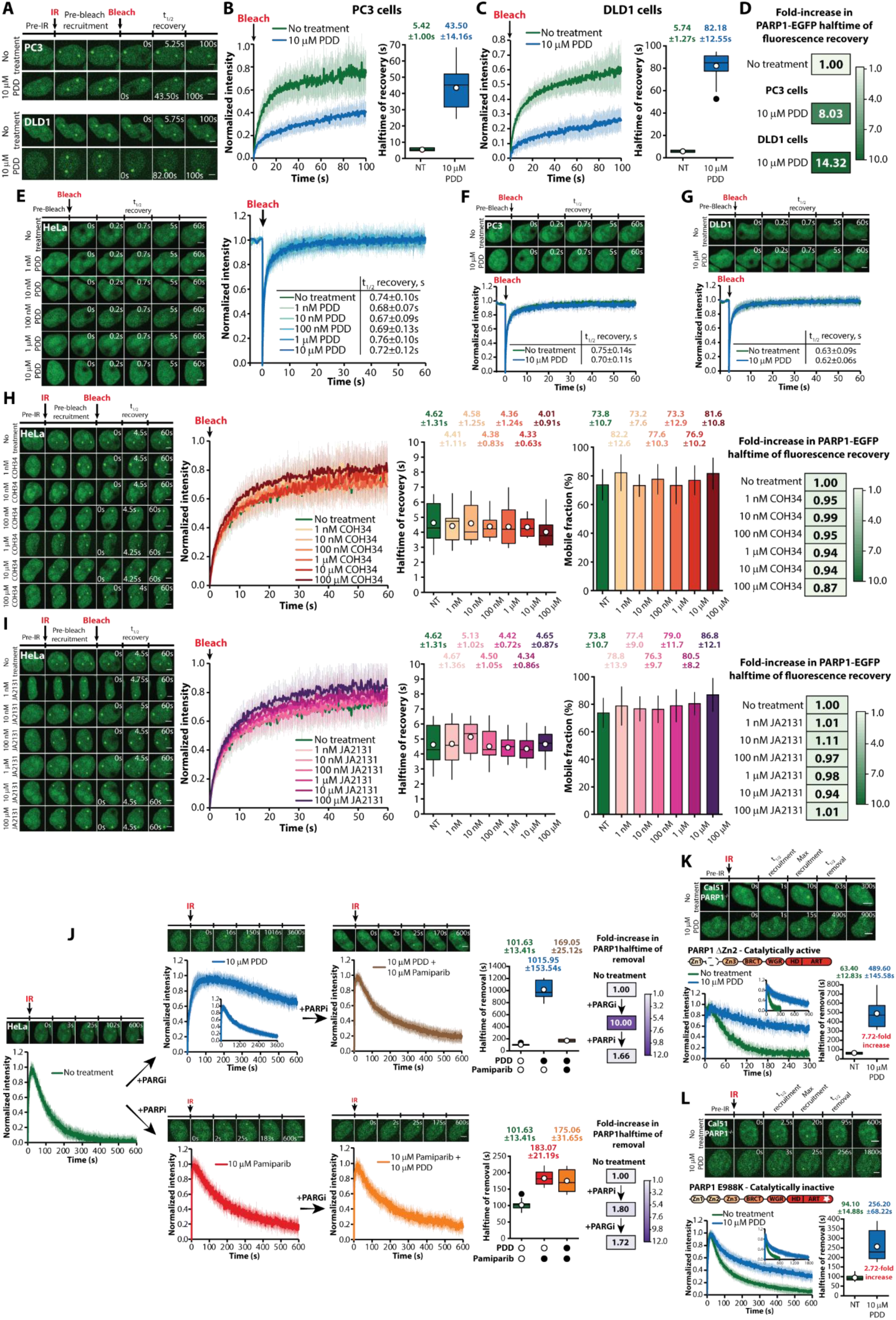
PARG inhibition slows down PARP1 turnover at damaged chromatin sites. (A-D) Time-lapse images, normalized kinetics, and fluorescence recovery half-times (including fold-increases) of PARP1 at IR-induced damage sites, as determined via FRAP in PC3 and DLD1 cells treated with PDD. PARP1-EGFP fluorescence recovery at non-damage sites, as determined via FRAP, in PDD-treated HeLa (E), PC3 (F), and DLD1 (G) cells. (H, I)Time-lapse images normalized kinetics, half-times of fluorescence recovery (including fold-increases), and mobile fraction (%) of PARP1 at IR-induced DNA damage sites as determined via FRAP in HeLa cells treated with a range of concentrations of COH34 and JA2131, respectively. (J) Time-lapse images, normalized kinetics, and PARP1 removal half-times (including fold-increases) in HeLa cells sequentially treated with PDD and pamipariborvice versa. (K, L) Time-lapse images, normalized kinetics, and removal half-times of AZn2 and E988K PARP1 transiently expressed in Cal51 PARP1-/- cells treated with PDD. Data are presented as the mean±SD. White dots indicate the mean value. Scale bars: 5 pm. NT, no treatment; IR, UV laser micro-irradiation.

**Figure S3.**
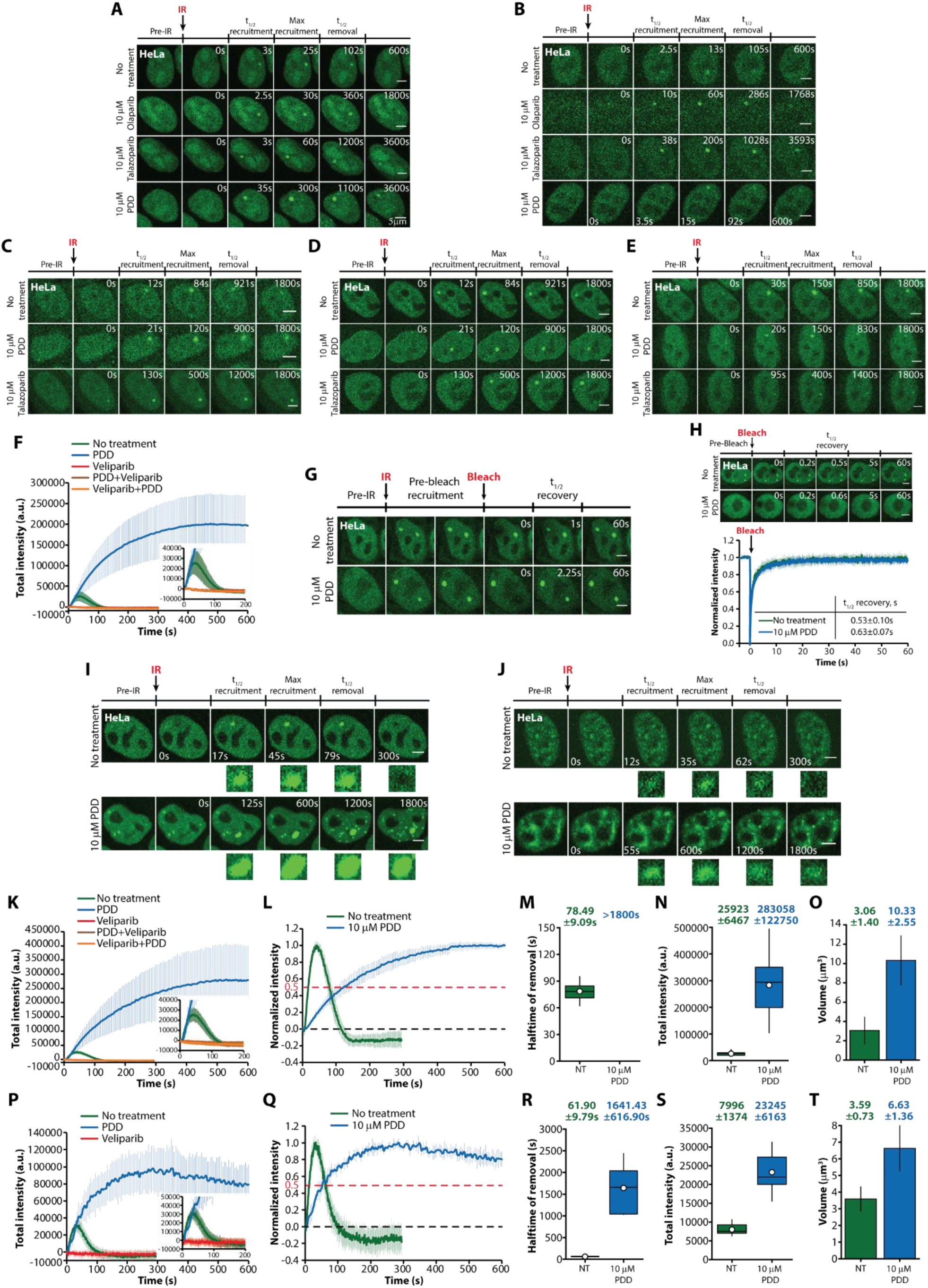
Effects of PARG inhibition on downstream repair factor and FET family protein dynamics. (A, B) Time-lapse images of PARP1 (A) and Timeless (B) at IR-induced damage sites following treatment with olaparib, talazoparib, or PDD. (C-E) Time-lapse images of RFC4 (C), PCNA (D), and POLD2 (E) at IR sites following treatment with talazoparib or PDD. (F) Total intensity kinetics of FUS at IR sites following treatment with PDD, veliparib, or both. (G) Time-lapse images of FRAP of FUS foci at IR-induced damage sites. (H) Time-lapse images and normalized intensity of FUS fluorescence recovery after photobleaching of a nuclear region that was not subjected to IR. Half-times of fluorescence recovery are also indicated. (I, J) Time-lapse images of EWSR1 (I) andTAF15 (J) recruitment and removal from IR damage sites, with and without PDD. (K-O) Total intensity kinetics (K, N), normalized kinetics (L), half-times of removal (M), and volume (O) of EWSR1 IR foci after treatment with PDD, veliparib, or both. (P-T) Total intensity kinetics (P, S), normalized kinetics (Q), half-times of removal (R), and volume (T) ofTAF15 IR foci after treatment with PDD or veliparib. Data are presented as mean±SD. White dots indicate the mean value. Scale bars: 5 pm. NT, no treatment; IR, UV laser micro-irradiation; a.u., arbitrary units.

**Figure S4.**
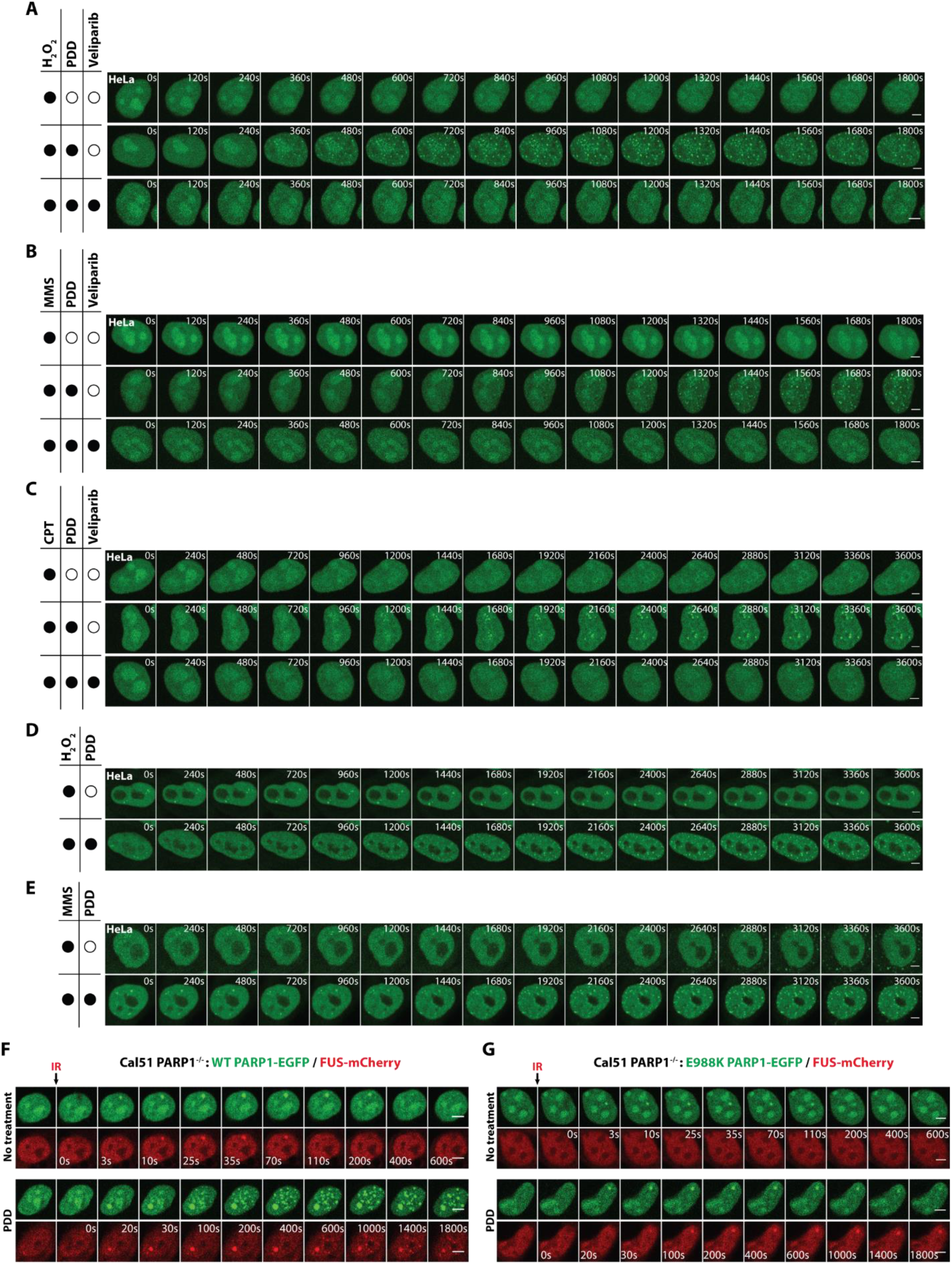
PARG inhibition induces aberrant PARP1 condensation in response to DNA damage. (A-C) Time-lapse images of PARP1 condensate formation in response to 100 pM hydrogen peroxide (H2O2) (A), 0.01% methyl methanesulfonate (MMS) (B), and 10 pM camptothecin (CPT) (C) treatment in HeLa cells pre-treated with 10 pM PDD or 10 pM PDD + 10 pM veliparib. (D, E) Time-lapse images of FUS condensate formation in response to 100 pM H2O2 (D) and 0.01% MMS (E) treatment in HeLa cells pre-treated with 10pM PDD. (F) Time-lapse images of PARP1 and FUS condensate formation in response to IR in PDD-treated Cal51 PARP1-/- cells transiently co-expressing WT PARP1 and WT FUS. (G)Time-lapse images showing compromised PARP1 and FUS condensate formation in response to IR in PDD-treated Cal51 PARP1-/- cells transiently co-expressing the inactive E988K PARP1 mutant and WT FUS. Scale bars: 5 pm. IR, UV laser micro-irradiation.

**Figure S5.**
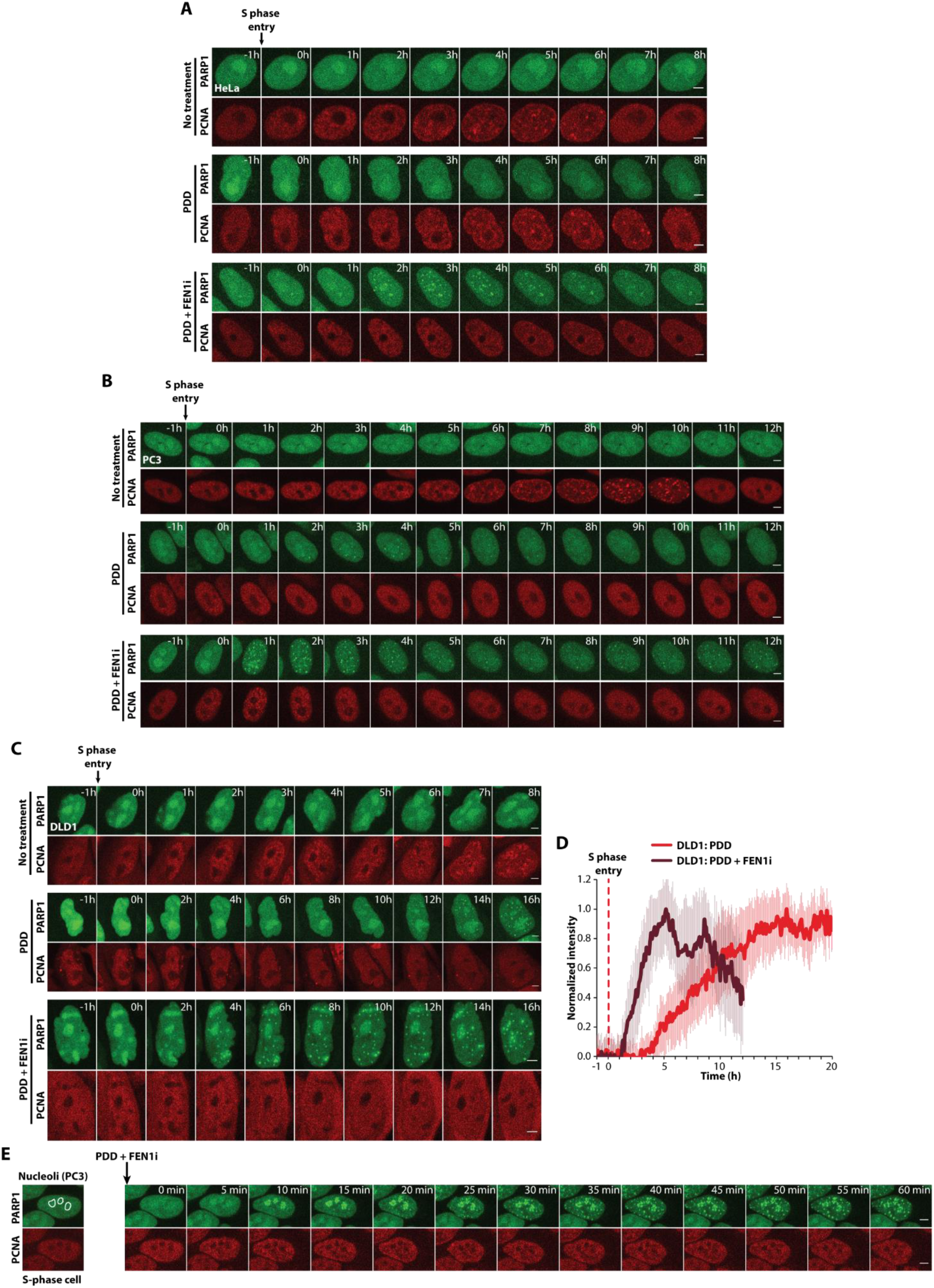
S-phase PARylation triggers PARP1 condensation. (A) Time-lapse images of HeLa cells followed throughout the cell cycle after S phase entry without treatment, with 10 μM PDD, or with 10 μM PDD + 10 μM FEN1-IN-1 (FEN1i). (B) Time-lapse images of PC3 cells followed throughout the cells cycle after S phase entry without treatment, with 10 μM PDD, or with 10 μM PDD + 10 μM FEN1i. (C) Time-lapse images of DLD1 cells followed throughout the cell cycle after S phase entry without treatment, with 10 μM PDD, or with 10 μM PDD + 10 μM FEN1i. (D) Normalized kinetics of formation of PARP1 foci in DLD1 cells treated with 10 μM PDD alone or in combination with 10 μM FEN1i. Data are presented as the mean+SD. Scale bars: 5μm.

**Figure S6.**
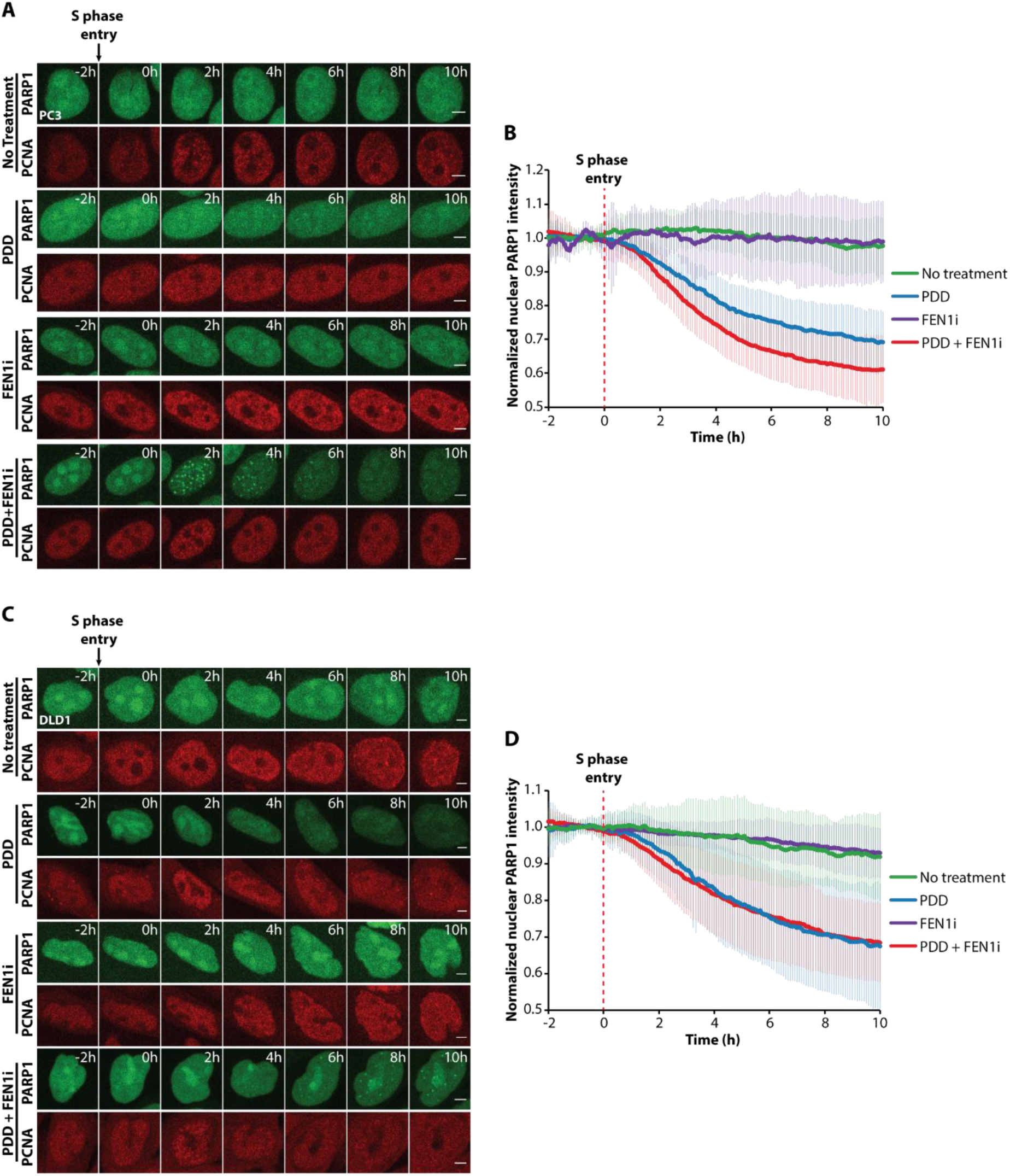
PARG-mediated dePARylation prevents the nuclear export of PARP1 during early S phase. (A, C) Time-lapse images of PARP1 -EGFP/ PCNA-mCherry-expressing PC3 (A) and DLD1 (C) cells progressing through S phase under treatment with 10 pM PDD, 10 pM FEN1-IN-1 (FENIi), or both.(B, D) Nuclear PARP1 fluorescence intensity in PC3 (B) and DLD1 (D) cells after treatment with 10 pM PDD, 10 pM FENIi, or both, normalized to the average intensity of 15 frames before S phase entry. Data are presented as the mean±SD. Scale bars: 5 pm.

